# Alprazolam modulates persistence energy during emotion processing in first-degree relatives of individuals with schizophrenia: a network control study

**DOI:** 10.1101/2021.04.22.440935

**Authors:** Arun S. Mahadevan, Eli J. Cornblath, David M. Lydon-Staley, Dale Zhou, Linden Parkes, Bart Larsen, Azeez Adebimpe, Ari E. Kahn, Ruben C. Gur, Raquel E. Gur, Theodore D. Satterthwaite, Daniel H. Wolf, Danielle S. Bassett

## Abstract

Schizophrenia is marked by deficits in facial affect processing associated with abnormalities in GABAergic circuitry, deficits also found in first-degree relatives. Facial affect processing involves a distributed network of brain regions including limbic regions like amygdala and visual processing areas like fusiform cortex. Pharmacological modulation of GABAergic circuitry using benzodiazepines like alprazolam can be useful for studying this facial affect processing network and associated GABAergic abnormalities in schizophrenia. Here, we use pharmacological modulation and computational modeling to study the contribution of GABAergic abnormalities toward emotion processing deficits in schizophrenia. Specifically, we apply principles from network control theory to model persistence energy – the control energy required to maintain brain activation states – during emotion identification and recall tasks, with and without administration of alprazolam, in a sample of first-degree relatives and healthy controls. Here, persistence energy quantifies the magnitude of theoretical external inputs during the task. We find that alprazolam increases persistence energy in relatives but not in controls during threatening face processing, suggesting a compensatory mechanism given the relative absence of behavioral abnormalities in this sample of unaffected relatives. Further, we demonstrate that regions in the fusiform and occipital cortices are important for facilitating state transitions during facial affect processing. Finally, we uncover spatial relationships (i) between regional variation in differential control energy (alprazolam *versus* placebo) and (ii) both serotonin and dopamine neurotransmitter systems, indicating that alprazolam may exert its effects by altering neuromodulatory systems. Together, these findings reveal differences in emotion-processing circuitry associated with genetic vulnerability to schizophrenia.

## Introduction

Schizophrenia is associated with deficits in emotion processing. Individuals with schizophrenia demonstrate marked deficits in facial affect perception, as measured through tasks that require the identification of emotions such as happiness, sadness, anger or fear^1,2^. Emotion processing deficits in schizophrenia contribute substantially to impairments in social cognition and poor functional outcomes^3,4^. First-degree family members of individuals with schizophrenia also display abnormalities in facial affect perception, albeit to a lesser extent than probands^5–8^. Abnormalities in first-degree relatives are particularly remarkable as the study of family members allows for the investigation of schizophrenia associated endophenotypes without the confounding effects of antipsychotic medication and secondary effects related to disease chronicity^9^. More broadly, investigations of facial affect processing in family members may offer insight into a key cognitive domain adversely affected by schizophrenia and can serve to inform effective treatment strategies.

Prior studies have used neuroimaging to characterize the neural circuitry associated with altered facial affect processing in individuals with schizophrenia and their relatives. These studies have primarily focused on linking differences in activation of limbic regions like the amygdala with altered identification and recall of threat-related faces^9–12^. However, facial affect perception is a complex process involving multiple brain regions, and evidence exists for impairment in both emotion-processing limbic regions as well as early-stage visual processing in schizophrenia^9,13^. Facial affect processing involves a distributed network comprising limbic regions, fusiform and occipital cortex, medial and lateral prefrontal areas, and insula^14–16^. Indeed, components of this distributed network have been implicated in facial emotion processing abnormalities in individuals with schizophrenia^17^ and individuals with high genetic risk for schizophrenia^18^, suggesting heritability. Thus, an integrative understanding of facial affect processing abnormalities in schizophrenia requires analysis of the distributed network regulating a complex domain.

Facial affect processing abnormalities in schizophrenia, and other cognitive deficits, may be driven by abnormal GABAergic neurotransmission^19^. Notably, GABAergic abnormalities in schizophrenia have been documented quite broadly, across the prefrontal cortex^20^, visual cortex^21,22^, amygdala^23^, and temporal lobe^24^, regions that overlap with the distributed network involved in facial affect processing. The role of GABAergic circuitry in facial affect processing and its impairment in schizophrenia can be effectively studied through pharmacological modulation using GABA modulators like benzodiazepines^25^. Alprazolam (Xanax®) is among the most widely used benzodiazepines, with well-known anxiolytic effects through enhanced GABAergic inhibition of the amygdala and limbic structures, and sedative effects from more broad GABAergic inhibition^26–28^. Thus, benzodiazepine challenge provides an opportunity to study the role of GABAergic circuitry in the etiology of facial affect processing abnormalities in schizophrenia.

Benzodiazepines impair emotion identification and emotion memory in healthy individuals, mainly in processing threatening faces^29–31^. The neural basis for the observed impairments in threat processing have been investigated in neuroimaging studies of emotion processing with benzodiazepine challenge in healthy subjects. These studies have shown that benzodiazepines alter activation of brain regions in the distributed facial affect processing network including amygdala, fusiform gyrus, orbitofrontal cortex, and insula during facial affect processing tasks^32,33^. We showed that alprazolam unmasks amygdalar and/or GABAergic abnormalities in first-degree relatives of individuals with schizophrenia during emotion identification and recall tasks^34^. However, there remains a lack of mechanistic understanding of benzodiazepine action during facial affect processing that goes beyond traditional activation studies. More recent tools for modeling the dynamics of brain activation states can help to synthesize results from activation studies and provide mechanistic insight into benzodiazepine action as well as GABAergic abnormalities in schizophrenia.

The mechanistic basis of benzodiazepine action on the brain during emotion processing can be effectively modeled using network control theory (NCT). NCT is a tool originating in theoretical physics and systems engineering that has successfully been used to understand how to control real-world systems comprised of interacting components, such as power grids and electronic circuits^35,36^. In the context of NCT, control refers to the ability to drive the system, through a suitable choice of inputs, from an initial state to a final state. Given that the brain is a complex system comprised of interconnected networks of neurons^37^, NCT provides an intuitive and compelling tool to model the dynamic trajectory of brain activation states that support its rich cognitive functions. Indeed, NCT has already been used to provide insight into the structure and function of model nervous systems like *C. elegans*^38^, *Drosophila*^39^, mouse^39,40^, and macaque^41^, as well as human brain networks^39,42–46^.

The application of NCT to model the brain typically involves the definition of a structural network through diffusion weighted imaging, and the definition of brain states as activation patterns across brain regions^47^. Brain states can be defined by arbitrarily switching ‘on’ canonical brain sub-networks like the visual and default mode networks, or directly as task activation obtained through functional magnetic resonance imaging data^43,48–51^. The NCT framework is then used to model the temporal progression of brain states as a function of the underlying structural network and theoretical control energy applied to different brain regions. The calculated control energy may represent external electrical stimulation or internal cognitive control needed to steer the brain between defined initial and final states^42^. Additionally, the brain regions important for driving specific brain state transitions can be identified through control impact analysis. This framework naturally lends itself to modeling the effect of drugs like alprazolam in driving brain state trajectories relevant to facial affect processing and can provide mechanistic insight into the mode of action of the drug beyond simple measures of activation.

Here we applied principles from network control theory to investigate the effect of alprazolam and schizophrenia risk status in driving brain state transitions during facial affect processing. We leveraged a previously reported dataset^34^ where fMRI BOLD data was collected during emotion identification and emotion memory tasks, with and without administration of alprazolam, in a cohort consisting of healthy controls and unaffected first-degree relatives of individuals with schizophrenia. We considered task-evoked brain activation patterns during emotion processing tasks to be brain states and quantified the theoretical control energy needed to maintain those states – the persistence energy. In our previous study, we showed that alprazolam unmasked GABAergic abnormalities in the amygdala in relatives^34^. Accordingly, our primary hypothesis was that when administered alprazolam, family members would have altered persistence energy during identification and recall of threatening faces which requires amygdalar processing, but not during non-threatening or neutral stimuli. We predicted that brain regions of high control impact in the NCT model would align with known regions involved in facial affect processing including fusiform cortex, occipital cortex, and sub-cortical regions like the amygdala and insula. Finally, we predicted that regions of high control impact would also spatially align with regions of high GABA receptor expression, but not with other neurotransmitters like dopamine and serotonin, reflecting the biological mode of benzodiazepene action. By testing and validating our hypotheses, we uncover novel insights regarding the contribution of GABAergic abnormalities toward emotion processing deficits in schizophrenia.

## Methods

### Participants

The sample included 27 healthy participants with a first-degree relative affected by schizophrenia and 20 healthy controls without a family history of schizophrenia, for a total of n=47 participants. Controls and relatives were matched based on demographic and clinical variables (**Table 1**). After excluding scans based on motion estimates (mean framewise displacement > 0.5mm), the final sample for data analysis included n=44 participants (19 relatives; 25 controls) for emotion identification, and n=40 participants (17 relatives; 23 controls) for emotion memory (see Supplementary Methods for details on assessment). Study procedures were approved by the University of Pennsylvania Institutional Review Board, and written informed consent was obtained from participants. Participants underwent standard medical, neurological, psychiatric, and neurocognitive evaluations (see Supplementary Methods).

**Table 1.** Demographic and clinical information at time of scan.

| Variable | Controls (n=27) |  | Relatives (n=20) |  | *p*-value | Odds ratio |
| --- | --- | --- | --- | --- | --- | --- |
|  | Percentage | Proportion | Percentage | Proportion |  |  |
| Sex (% F) | 51.9 | 14F/13M | 55.0 | 11F/9M | 1.0 | 0.88 |
| Handedness (% R) | 92.6 | 25R/2L | 80.0 | 16R/4L | 0.38 | 0.32 |
| Smoke (% N) | 77.8 | 21N/6Y | 80.0 | 16N/4Y | 1.0 | 0.88 |
|  | Mean (SD) | Range | Mean (SD) | Range |  | Test statistic* (DOF) |
| Age (years) | 39.0 (11.4) | 21.1-56.5 | 42.3 (14.8) | 20.9-59.4 | 0.31 | -1.02 |
| Education (years) | 15.0 (2.0) | 11.0-19.0 | 14.8 (2.3) | 12.0-20.0 | 0.79 | 0.26 (45) |
| Parental education | 13.6 (3.1) | 7.5-20.0 | 13.9 (2.7) | 9.5-18.0 | 0.76 | -0.31 (43) |
| Height (in.) | 67.7 (4.0) | 61.0-77.0 | 67.6 (4.3) | 60.0-73.0 | 0.93 | 0.09 (45) |
| Weight (lb.) | 176.4 (33.0) | 115.0-255.0 | 175.5 (34.0) | 118.0-250.0 | 0.93 | 0.09 (45) |
| BMI (lb./in.<sup>2</sup>) | 27.1 (4.9) | 20.4-36.8 | 27.0 (4.5) | 18.7-33.9 | 0.94 | 0.08 (45) |
| Trait anxiety | 28.3 (6.7) | 20.0-47.5 | 30.1 (8.9) | 20.0-58.0 | 0.64 | -0.46 |
| Schizotypy (SIS) total | 11.5 (7.2) | 1.0-29.0 | 15.0 (7.2) | 7.0-39.0 | 0.11 | -1.64 (44) |
| Alprazolam level (ng/mL) | 7.5 (4.1) | 0.0-13.0 | 7.8 (4.1) | 1.0-14.0 | 0.80 | -0.26 (42) |

Notes: F=female, M=male, R=right, L=left, N=non-smoker, Y=smoker, SD=standard deviation, BMI=body mass index, SIS=structured interview for schizotypy, DOF = degrees of freedom; reported *p*-values are from Fisher's exact test for categorical variables (sex, handedness, and smoking status), Wilcoxon rank sum tests for non-normal data (age, trait anxiety), and two-sample *t*-tests for normally distributed data (all other variables). \* *t*-statistic for two-sample *t*-tests, *z*-statistic for Wilcoxon rank sum test (degrees of freedom not reported).

### Study design and pharmacological challenge

To study the impact of GABAergic modulation on brain activation during emotion processing, participants underwent fMRI imaging during facial affect processing tasks with and without administration of alprazolam. Details of study design have been described previously^34^. Briefly, participants underwent two identical fMRI sessions approximately one week apart. Participants were administered 1mg oral alprazolam in one session and an identical-appearing placebo in the other session, in a balanced double-blind within-subject crossover design.

During each fMRI session, participants performed an emotion identification task followed by an emotion memory task. In the emotion identification task, 60 unique color pictures of human faces were presented in pseudorandomized order, with facial expressions falling into one of five emotional categories: happy, sad, fearful, angry or neutral^52^. Participants were asked to identify the emotion expressed on each face. In the emotion memory task, the same sequence of faces as in the preceding emotion identification task was presented, with each target face accompanied by two foil expressions. Participants were instructed to recall the expression that matched the previously seen face. In both tasks, each emotion category was presented 12 times, with each emotion being used as a foil 24 times in the emotion memory task. Faces were displayed for 5.5s, with a variable interval of between 0.5-18.5s, during which a complex crosshair matched to faces on perceptual qualities was presented. Each task lasted 10.5 min, with a 2 min delay between tasks.

### Image acquisition and processing

Structural and functional image sequences were acquired with a Siemens Trio 3T system (Erlangen, Germany). Structural images were acquired for the whole brain, whereas functional volumes were acquired in a slab covering ventral regions of the brain with a spatial resolution of 2×2×2 mm (Figure S1).

We used *fMRIPrep* software (version 1.2.6) to process the BOLD fMRI data^53^. Briefly, *fMRIPrep* was used to perform brain extraction and segmentation of individual T1-weighted images, spatial normalization of T1 images to the ICBM 152 Nonlinear Asymmetrical template, susceptibility distortion correction for BOLD images, estimation of confound variables including head motion parameters and resampling of BOLD sequences to MNI152NLin2009cAsym standard space. We excluded sessions for which the average framewise displacement was greater than 0.5mm. No other exclusion criteria were applied.

Next, we used generalized linear models (GLM) to measure subject-specific brain activation patterns during emotion identification and memory tasks. Specifically, GLM analysis was performed using the FEAT module^54^ in FSL 5.0.10 implemented using XCP Engine^55^. BOLD sequences preprocessed using *fMRIPrep* were high-pass filtered (100s) and spatially smoothed (4mm FWHM, isotropic); further, the first 6 non-steady state volumes were discarded. All event conditions were modeled as 5.5s-boxcars convolved with a canonical hemodynamic response function. Consistent with previous work^15,56^, correct responses to fear and anger stimuli were combined as a “threat” regressor; happy and sad stimuli were combined as a “non-threat” regressor; and neutral stimuli were modeled separately. All specified contrasts measured BOLD activation compared to baseline. Incorrect responses and 6 motion parameters were included as regressors of non-interest. We chose to include only correct responses in the model to limit the potential effects of inattention due to sedation by alprazolam.

We then divided the brain into 233 parcels based on the Lausanne parcellation (after excluding the brain stem), which provides coverage of both cortical and subcortical areas including thalamus, caudate, putamen, pallidum, accumbens, hippocampus, and amygdala^57^. Parameter estimates (beta weights) from each voxel were averaged within each parcel resulting in estimates of brain activation (brain states); these activation maps were then evaluated using network control theory.

See Supplementary Methods for further details on image acquisition and processing.

### Construction of structural brain networks from diffusion spectrum imaging data

Structural brain connections are an essential component of network control theory models. Since we did not collect structural brain images in our previous study, we leveraged an average structural matrix from a separate study. Diffusion spectrum imaging (DSI) data was collected from a separate set of 10 healthy young adults as described elsewhere^47^. Consistent with previous work^48,49^, we defined nodes of the structural network as brain regions according to the Lausanne atlas^57^. To encode each structural network, we constructed adjacency matrices for each subject based on the quantitative anisotropy (QA) between each pair of brain regions. The average structural matrix across 10 participants was used for all results shown in the main text.

See Supplementary Methods for further details on DSI image acquisition, processing, and structural matrix generation.

### Network control theory

We used principles from network control theory (NCT) to investigate the effect of alprazolam and schizophrenia risk status in driving brain state trajectories associated with facial affect processing. The NCT framework has been used to determine how underlying white matter architecture constrains transitions between different brain states inferred from neuroimaging data^43,48–51^. We begin by approximating brain state dynamics through the linear continuous-time equation

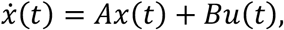

Where *x*(*t*) is a vector of size *N* × 1 (where *N* is the number of brain regions in the network) that represents the state of the system at time *t, A* is the weighted symmetric *N* × *N* structural matrix estimated through diffusion spectrum imaging, *B* is an input matrix of size *N* × *N* specifying the set of control nodes, and *u*(*t*) is the time-dependent control signal in each of the control nodes.

The minimal control energy framework^47^ defines the unique control input *u*\*(*t*) needed to transition the system from an initial state x(0) = *x*_0_ to a final target state *x*(*T*) = *x*_*T*_ over the time horizon *T* through the cost function

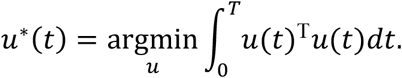

By integrating each control input over time, we can calculate the control energy required by each brain region as

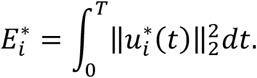

By summing the control inputs over all *N* brain regions, we can obtain the total control energy needed for the transition from *x*_0_ to *x*_*T*_, which we write as

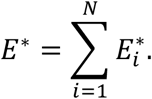

In our model, we used parameter estimates (beta weights from a general linear model) to specify brain states *x* from alprazolam and placebo sessions, and a single group-representative structural matrix *A* (**Figure 1**). Since BOLD images were acquired in a slab covering ventral regions, we restricted our analysis to parcels with at least 50% coverage within the slab (Figure S1). Thus, *x A*, and *B* matrices were truncated for each individual based on slab coverage. For simplicity, all nodes within the slab were set as controllers, thus allowing us to evaluate multi-point (as opposed to single-point) control.

**Figure 1.**
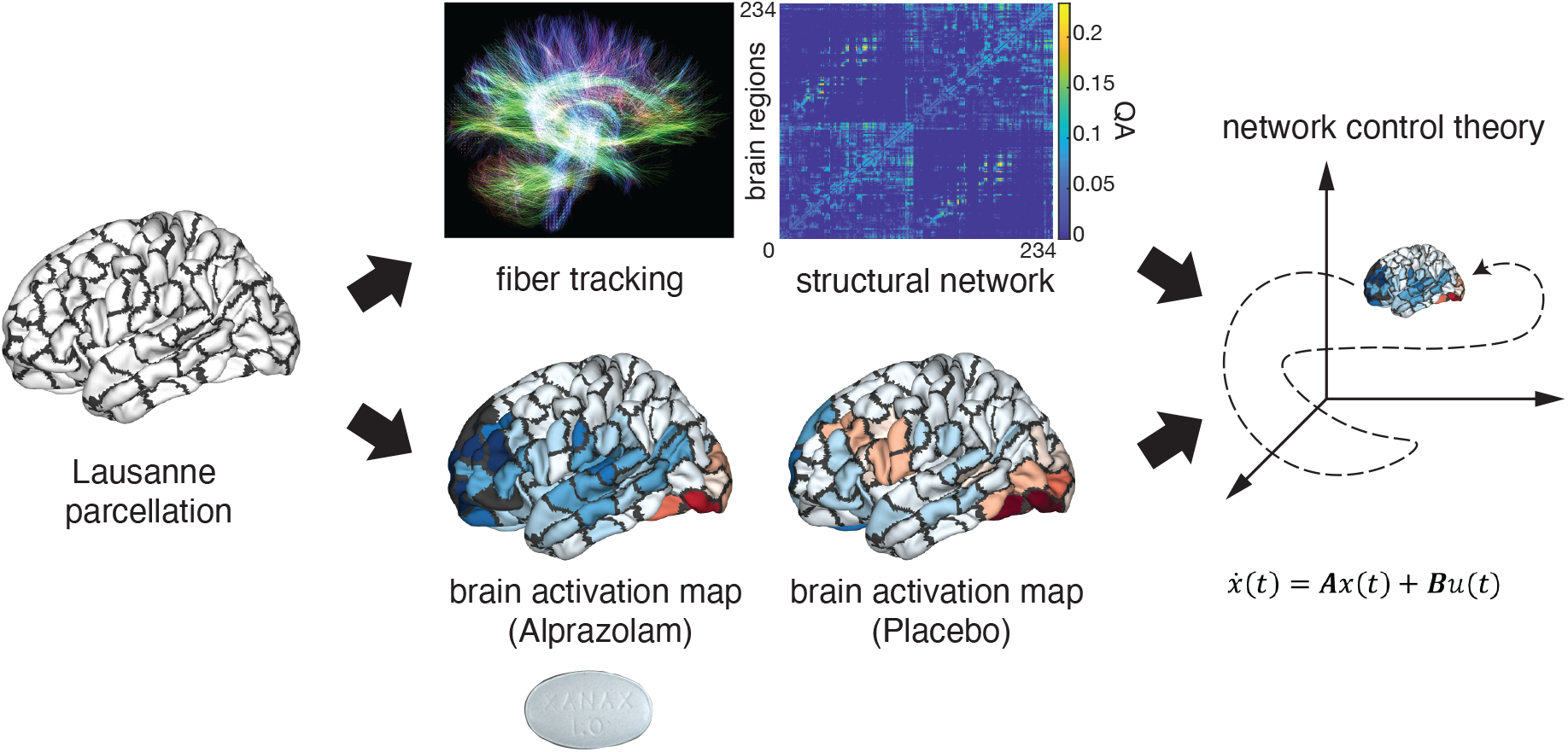
Operationalizing network control theory in the context of human neuroimaging. The strength of structural connections between brain regions were determined by the quantitative anisotropy (QA) estimated from diffusion spectrum imaging data. We used beta coefficients from general linear models to specify brain activation maps during task sessions where participants were given 1mg oral alprazolam or placebo. These maps were then fed into a network control model to analyze the energy required for transitions between different brain states. We were particularly interested to estimate the persistence energy,***P***_***e***_ defined as the energy required to maintain a state. Brain regions in the cortex and subcortex were defined by the 234-node Lausanne parcellation.

The impact of different modeling choices for the time horizon parameter *T* (time during which the control input is effective) was explored by calculating the Pearson correlation between persistence energies calculated for pairs of parameter values^47^. We found that short time horizons resulted in a different control regime as expected^47^; however, correlations were >0.99 overall, indicating that the choice of time horizon did not significantly affect minimal control energy calculations (Figure S2). Based on these findings, we chose a time horizon of *T* = 3 for our simulations.

We wished to extract control energy parameters that reflected the cognitive process of facial affect processing. Here, we used the notion of persistence energy, defined as the control energy needed to maintain a brain state^43,51^. For an initial brain state x_0_, we defined persistence energy P_e_ as the minimum control energy needed for the system to reach a final state x_T_ = x_0_, such that the initial state is maintained at time *T* (**Figure 1**). Persistence energy associated with different brain states can be interpreted as the cognitive effort exerted during the performance of tasks relevant to sustain those states^42,43^. States with larger overall magnitude are always more difficult to maintain; thus, to facilitate comparisons between persistence energies associated with different brain states independent of the global activation magnitude of each state, we divided each brain state vector, x, by its Euclidean norm.

In order to evaluate the importance of different brain regions in driving brain state transitions during facial affect processing, we measured the control impact *I*_*i*_ of individual nodes by iteratively removing each node from the network and recomputing the persistence energy^49^. Brain regions with highest control impact are those whose removal from the control set leads to the highest increase in control energy. Formally, the control impact is defined as

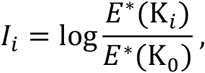

Where K_0_ represents the set of all control nodes and K_*i*_ represents the set of control nodes after excluding node *i*

### Spatial correlations with neurotransmitter maps

In order to explore the underlying biology of drug action reflected through control energy measures, we analyzed the spatial alignment of drug-induced differences in control energy input with neurotransmitter receptor maps obtained through PET imaging. Given the known role of alprazolam as a GABA modulator^27^, we expected brain regions whose control energy input varied strongly with drug condition to also be correlated with GABA receptor density, but not with other receptors such as serotonin and dopamine.

For this analysis, we used published PET/SPECT maps of the following receptors: 5-HT1a (serotonin 5-hydroxytryptamine receptor subtype 1a), 5-HT1b (5-HT subtype 1b), 5-HT2a (5-HT subtype 2a), D1 (dopamine D1), D2 (dopamine D2), DAT (dopamine transporter), F-DOPA (dopamine synthesis capacity), GABA_A_ (gamma-aminobutyric acid A receptor), NAT (noradrenaline transporter), and SERT (serotonin transporter)^58–64^. All provided PET/SPECT maps were voxel-wise average group maps of variable numbers of healthy volunteers, linearly rescaled to a range of 0 to 100^58^. We further averaged the PET/SPECT maps voxel-wise for each Lausanne parcel to obtain 233×1vectors, each of which represented a spatial map of the distribution of a given neurotransmitter. We then evaluated correlations between all PET/SPECT maps and region-wise difference maps in control input between alprazolam and placebo sessions for all subjects.

### Statistical analyses

Primary hypotheses concerning the effects of drug (alprazolam vs. placebo) and group (control vs. family) on persistence energy were evaluated using linear mixed models to accommodate the repeated measures data (with performance across two sessions nested within 44 participants in emotion identification and 40 participants in emotion memory). Categorical indicators for drug were modeled as 0=alprazolam and 1=placebo, and the categorical group indicator was modeled as 0=control and 1=relative. We also evaluated mixed effects models by replacing the categorical drug indicator with alprazolam blood levels and replacing the categorical group indicator with schizotypy scores measured as the total score on the structured interview for schizotypy (Supplementary Information).

Separate models were evaluated for each emotion category (threat, non-threat, neutral) and task (emotion identification, emotion memory), resulting in 6 models. For each emotion category, formal models were constructed at level 1 as follows.

Level 1:

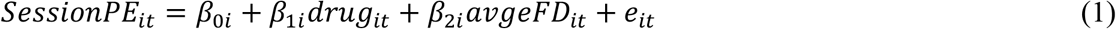

where *β*_0_ is the intercept, indicating the average level of persistence energy for the prototypical male control in an alprazolam session (determined by reference categories of categorical predictors); *β*_1*i*_ indicates within-person differences in persistence energy associated with drug or placebo sessions; *β*_2*i*_ indicates within-person differences in persistence energy associated with within-person differences in head motion for each session captured by average framewise displacement *(avgeFD_it_)* during each session, and *e*_*it*_ are autocorrelated session-specific residuals (AR1).

Person-specific intercepts and associations from level 1were specified at level 2 as follows.

Level 2:

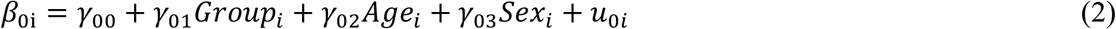

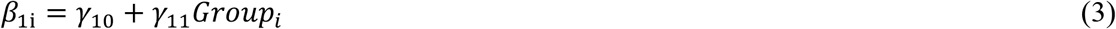

where *γ* denotes a sample-level parameter and *u* denotes residual between-person differences that may be correlated but are uncorrelated with *e*_*it*_ Parameters *γ*_01_) to *γ*_03_ indicate how between-person differences in the average persistence energy across sessions were associated with group, subject age, and subject sex. Parameter *γ*_11_ tests the moderating effect of group on the association between session persistence energy and drug administration. If there were no significant interactions, main effects were evaluated after removing interaction terms from the equations.

Exploratory analyses on relationships between persistence energy and task performance were conducted using a different set of linear mixed models. We evaluated separate models for each emotion category (threat, non-threat, neutral) and task (emotion identification, emotion memory), resulting in 6 models. Task performance measured as behavioral efficiency – proportion of correct responses divided by median reaction time for correct responses – was modeled as the dependent variable.

For each emotion category, formal models were constructed at level 1 as follows.

Level 1:

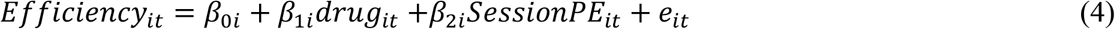

where *β*_0_ is the intercept, indicating the average task efficiency for the prototypical male control in an alprazolam session (determined by reference categories of categorical predictors); *β*_1i_ indicates within-person differences in task efficiency associated with drug or placebo sessions; *β*_2i_ indicates within-person differences in task efficiency associated with within-person differences in session persistence energy, and *e*_*it*_ are autocorrelated session-specific residuals (AR1).

Person-specific intercepts and associations from level 1were specified at level 2 as follows.

Level 2:

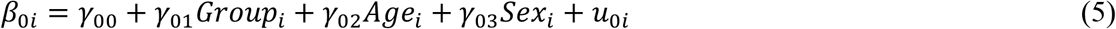

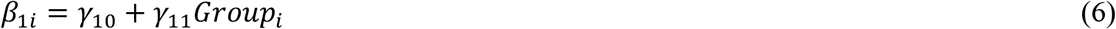

where *γ* denotes a sample-level parameter and *u* denotes residual between-person differences that may be correlated but are uncorrelated with *e*_*it*_ Parameters *γ*_01_ to *γ*_03_ indicate how between-person differences in task efficiency across sessions were associated with group, subject average persistence energy across sessions, subject age, and subject sex. Parameter *γ*_11_ tests the moderating effect of group on the association between task efficiency and drug administration. If there were no significant interactions, main effects were evaluated after removing interaction terms from the equations.

During exploratory analyses of spatial correlations between PET maps and region-wise difference maps in control input between alprazolam and placebo sessions, statistical significance was evaluated using permutation tests. Specifically, correlations were recomputed after randomizing PET spatial maps 10,000 times and *p*-values were estimated as the fraction of those iterations where the observed average correlation exceeded the randomized average correlation. Significant associations in all exploratory analyses were evaluated using the false discovery rate (FDR) to account for multiple comparisons^65^.

## Results

### Alprazolam differentially modulates persistence energy in relatives and controls during threat emotion processing

We first tested our primary hypothesis of alprazolam altering persistence energy during threat emotion processing. Persistence energy was measured as the control energy needed to maintain specific brain activation patterns observed during in-scanner emotion identification and memory tasks. We evaluated the effect of group and drug on persistence energy using linear mixed models with drug and group treated as categorical variables (Equations 1-3). During emotion identification, there was no main effect of group or drug, and no group×drug interaction in any emotion category (**Figure 2A**, see Supplementary Data Files 1 for model coefficients and statistics). During emotion memory, we found that persistence energy was significantly increased with alprazolam administration during recall of threat stimuli, in family members but not in controls (**Figure 2B**, group×drug interaction, *γ*_11_=-0.048, p=0.026, df=73). There was no main effect of group (*γ*_01_=0.026, p=0.133, df=73) or drug (*β*_1*i*_=0.01, p=0.50, df=74) (Supplementary Data Files 1). As expected, no significant effects were found in non-threat and neutral conditions. Alternate analyses where categorical drug indicator was replaced with alprazolam blood levels and categorical group indicator was replaced with the total score on the structured interview for schizotypy (SIS) showed similar results (Supplementary Information, Supplementary Data Files 2-3).

**Figure 2.**
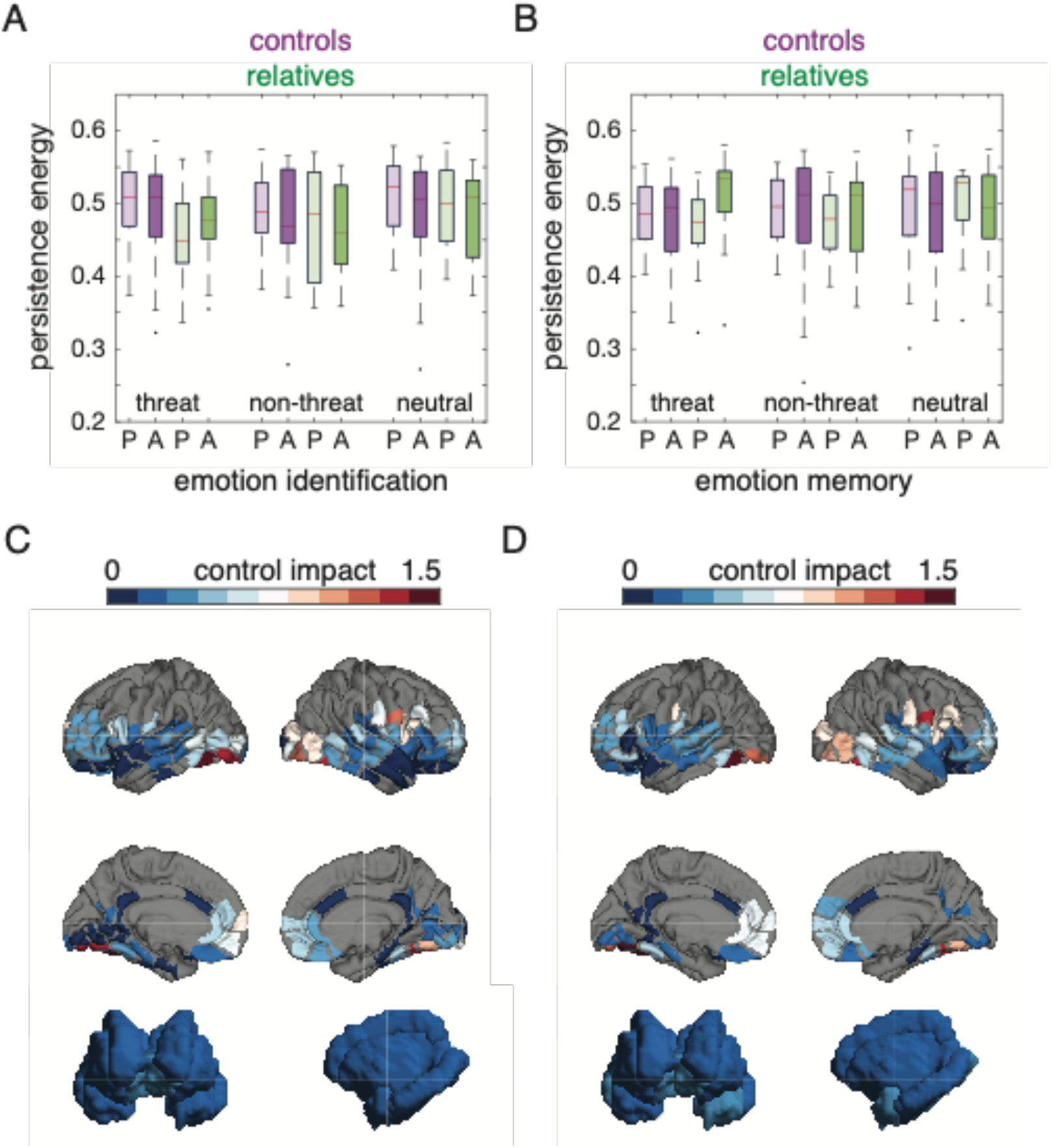
Alprazolam modulates persistence energy during recall of threatening faces. (A) Boxplots show persistence energy for the emotion identification task, grouped by emotion category; P=placebo; A=alprazolam. (B) Boxplots show persistence energy for emotion memory task, grouped by emotion category. We observed a significant group×drug interaction in the threat condition (***γ*_11_**=-0.048, p=0.026, df=73); P=placebo, A=alprazolam. (C) Average spatial maps of control impact for threat emotion identification, shown on surface renderings of cortical and subcortical areas. (D) Average spatial maps of control impact for threat emotion identification, shown on surface renderings of cortical and subcortical areas. Parcels outside the imaging slab are colored gray.

In order to elucidate the influence of structural brain networks and spatial activation patterns on the observed results, we performed a series of investigations using structural and spatial null models (see Supplementary Methods for details). These investigations showed that the differential effect of alprazolam on persistence energy in relatives and controls during recall of threatening faces was driven partially by structural brain networks but largely by spatial activation patterns (Figure S4, Figure S5, Supplementary Data Files 8, 9).

Finally, we used control impact analysis to investigate the relative importance of different brain regions in driving brain state transitions associated with emotion identification and memory. Control impact of individual nodes was measured by iteratively removing each node from the network and recomputing the persistence energy^49^.

As hypothesized, we found that regions with high control impact in both emotion identification and memory tasks were primarily located in the fusiform and occipital cortex, reflecting the visual and facial processing nature of the tasks (**Figure 2C-D**, Supplementary Data Files 4). One node in the precentral gyrus also exhibited high control impact, likely reflecting voluntary motor control for button presses during task execution. Surprisingly, subcortical areas including the amygdala, hippocampus, and insula had relatively low control impact. Areas with high control impact aligned largely with areas of high activation obtained from beta weight maps estimated from a general linear model (Figure S3, Supplementary Data Files 5). Thus, the control model suggests that the direct and indirect connectivity of the fusiform and occipital regions with the whole brain structural network facilitates efficient coordination of neural dynamics associated with facial affect processing.

### Individual differences in persistence energy explain variance in task performance during threat emotion identification

Next, we performed an exploratory analysis to examine the relationship between persistence energy and task performance. We reasoned that increased persistence energy during emotion identification and memory tasks might reflect cognitive effort expended and thus might be reflected in measures of task performance. We summarized task performance using an efficiency measure (accuracy divided by reaction time), and then evaluated associations between efficiency and persistence energy using a different set of linear mixed effects models (Equations 4-6).

We found that efficiency during threat emotion identification was positively associated with persistence energy (**Figure 3A**, main effect of persistence energy, β=0.335, p_FDR_=0.047, df=81). No significant associations were found for other emotion categories, or for the emotion memory task, after correction for multiple comparisons (**Figure 3B-F**). Consistent with our previous study^34^, we found that alprazolam significantly reduced task efficiency during both emotion identification and memory tasks (see Supplementary Data Files 6 for all model coefficients and statistics). We also found, consistent with our previous study, that there were no group effects on task performance for any of the task conditions (Supplementary Data Files 6). Thus, relatives and controls performed equally well on emotion identification and memory tasks.

**Figure 3.**
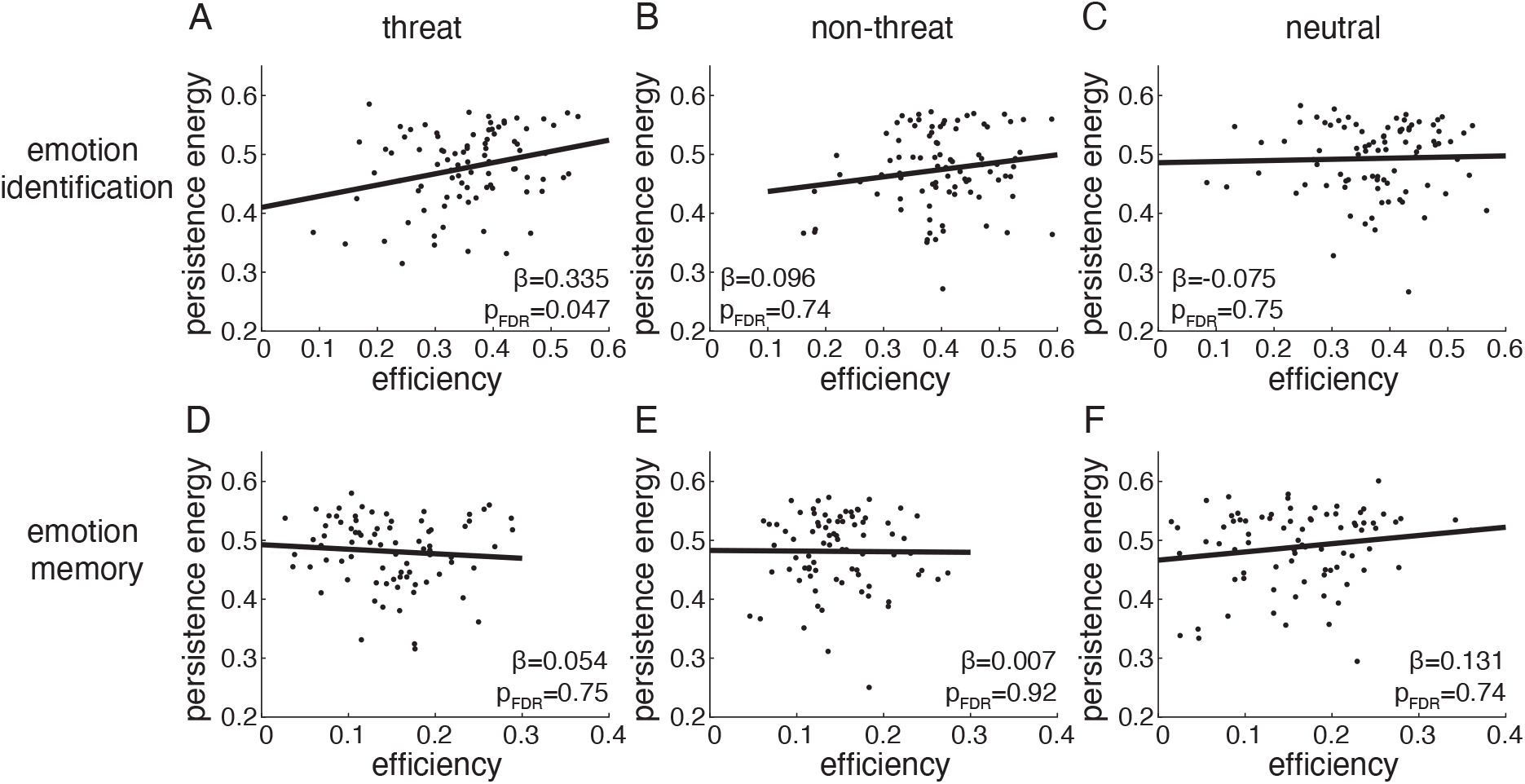
Task performance during threat emotion identification can be predicted from persistence energy. Scatterplots of efficiency in task performance against persistence energy, shown for the emotion identification (panels A-C) and emotion memory (panels D-F) tasks, separately for the threat, non-threat, and neutral categories. Task performance efficiency is measured as the proportion of correct responses divided by the median reaction time for correct responses. The ***β*** weights from the linear mixed effects models containing drug, group, age, and sex as covariates are shown on each plot, along with associated p-values corrected for multiple comparisons. Associations are significant for threat emotion identification at a significance level of p_FDR_<0.05.

### Regional differences in control energy spatially align with neurotransmitter systems

Finally, we explored the underlying biology of control energy measures by evaluating the spatial correspondence between control energy parameters from our model and known neurotransmitter systems described through PET/SPECT receptor maps^58^. To achieve this, we calculated the spatial correlation between PET/SPECT maps (**Figure 4A-B**) and maps describing regional differences in control energy input between alprazolam and placebo conditions (**Figure 4C**, Supplementary Data Files 7). Given the known role of alprazolam as a GABA_A_ receptor modulator^27^, we expected brain regions whose control input varied strongly with drug condition to also be correlated with GABA_A_ receptor density.

**Figure 4.**
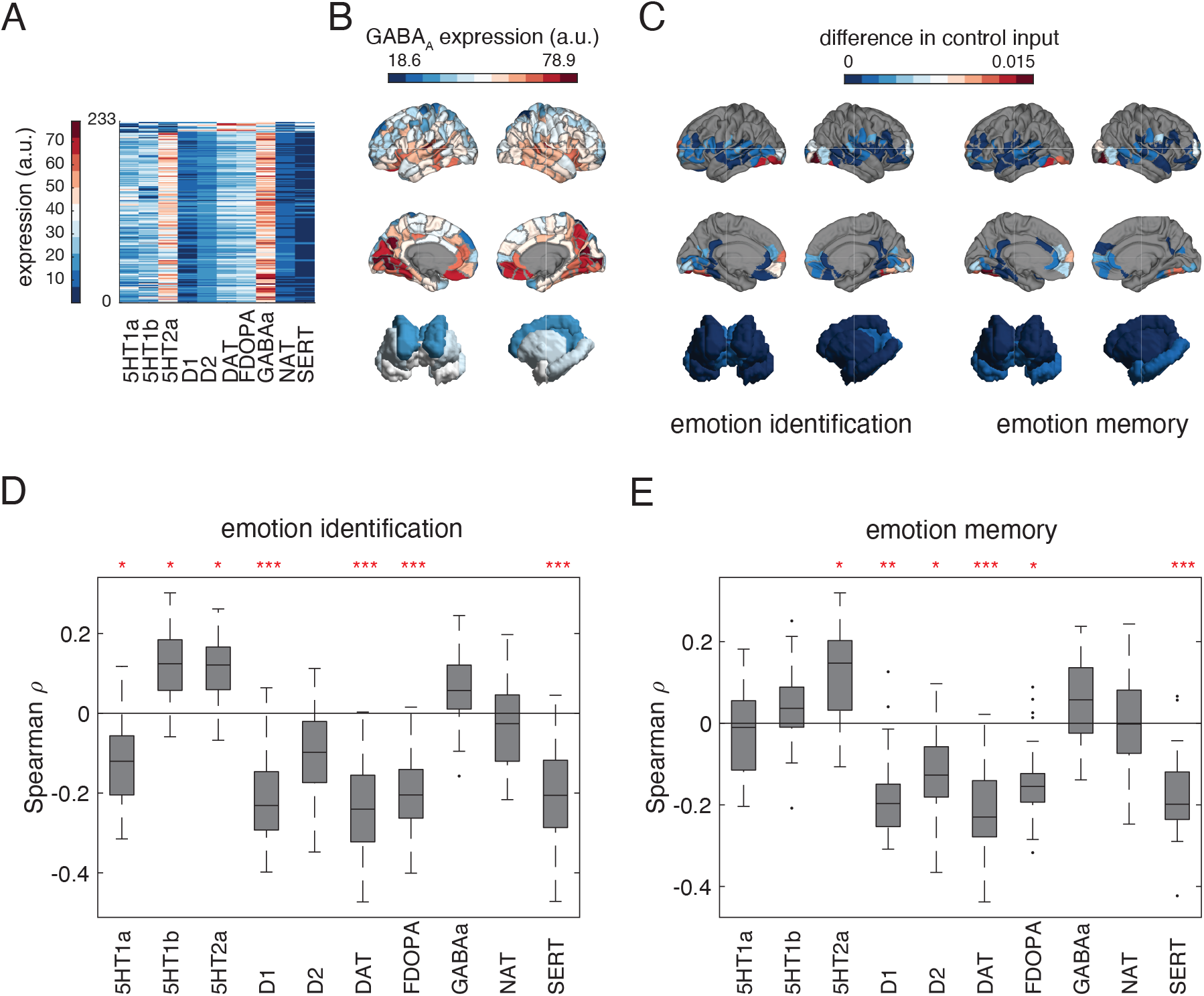
Neurotransmitter receptor profiles are associated with drug effect on control input. (A) PET neurotransmitter heatmaps from Dukart *et al*. (2020). (B) PET map of GABA_A_ expression shown on surface renderings of cortical and subcortical areas. (C) Regional differences in average control input on alprazolam and placebo (absolute values) during threat emotion identification and memory, shown on surface renderings of cortical and subcortical areas. (D-E) Boxplots of subject-level Spearman correlation coefficients between PET spatial maps and regional control input differences during threat emotion ID (panel D) and threat emotion memory (panel E). Red asterisks indicate the level of statistical significance from permutation tests with 500 permutations, corrected for multiple comparisons; * p_FDR_ < 0.05, ** p_FDR_ < 0.005, *** p_FDR_ < 0.0005.

To calculate regional control input difference maps, we subtracted total control input in each brain region (over the simulation time) between alprazolam and placebo sessions. These maps show that the effects of alprazolam are mainly located in occipital and fusiform areas, with some effects in frontal and orbitofrontal regions (**Figure 4C**, Supplementary Data Files 7). We then evaluated correlations between regional control input difference maps and PET/SPECT receptor maps. Surprisingly, we found that correlations between control input difference maps and GABA_A_ receptors were not significant (**Figure 4D-E)**. Moreover, in both emotion identification and memory tasks, control input difference maps were positively correlated with serotonergic receptors, and negatively correlated with dopaminergic receptors (**Figure 4D-E)**.

## Discussion

In this study, we applied a network control theory model to investigate the effects of alprazolam during facial affect processing in a cohort of healthy controls and first-degree relatives of people with schizophrenia. The main findings from our analysis and their implications are discussed below.

### Control energy measures as an endophenotype of schizophrenia

In our previous study^34^, we found that alprazolam effects on standard task fMRI measures in amygdala were stronger in relatives of individuals with schizophrenia compared to controls during emotion identification, suggesting alprazolam could be unmasking underlying GABAergic abnormalities. Given these prior results, we expected that alprazolam would also differentially influence control energy measures associated with whole-brain emotion-processing activation patterns in relatives versus controls. Indeed, we found that alprazolam increased the persistence energy associated with brain states during the recall of threatening faces (anger and fear) in family members but not in controls.

The persistence energy is the control energy needed to maintain a brain state associated with a task and has been previously associated with the cognitive effort required during those tasks^43^. We found further evidence for this relation between energy and effort by demonstrating that increased persistence energy is associated with better task performance in a subset of tasks. Since family members demonstrated relatively normal behavioral performance, increased persistence energy during the processing of threatening faces may represent a compensatory GABAergic mechanism that enables them to perform as well as controls. Further investigation into potential compensatory mechanisms might uncover promising avenues to target therapeutic drugs that aim to support improved cognitive function in schizophrenia.

Control energy measures go beyond traditional measures of activation, instead reflecting brain-wide network dynamics constrained by underlying white matter architecture^43,50,51^. Since facial affect processing is known to involve a distributed network of brain regions^14,15^, models that capture network-wide brain dynamics are important to investigate the neural substrate of this cognitive domain and its modulation by psychiatric disease. Our results add evidence of GABAergic abnormalities in family members when processing faces with negative affect, unmasked by drug action. Importantly, these abnormalities were measured using network-wide readouts, demonstrating that our analyses provide an important complementary approach to identifying such effects. Taken together, our results indicate that control energy measures could potentially be useful as an endophenotype of schizophrenia.

### Regions most impacting network control align with the facial affect processing network and distributions of neuromodulatory receptors

We sought to understand the impact of different brain regions in driving brain state transitions associated with emotion processing, expecting that regions of high importance would align with the distributed network associated with facial affect processing^14–16^. In partial support of this hypothesis, we found that brain regions with high control impact during emotion identification and memory were primarily in the fusiform and occipital cortices. Fusiform and visual brain regions are core components of the classical network associated with facial affect processing^14,15,66,67^. Our mathematical model suggests that the direct and indirect connectivity of these regions with the whole brain structural network facilitates efficient coordination of neural dynamics associated with performance of face processing behavior, providing novel intuition regarding their role as the “face areas” of the brain.

Our analysis also showed that limbic and sub-cortical regions such as amygdala, hippocampus and insula did not have high control impact in any emotion category. These regions have been classically associated with emotion processing^16,68^, and their low prominence in our network control model is therefore somewhat surprising. Our results may suggest that primary sensory areas associated with visual processing exert top-down control on whole-brain activation during facial affect processing, while subcortical regions including the amygdala are circumscribed to a bottom-up role with limited impact on the rest of the brain. Further, the high prominence of fusiform and occipital regions and low prominence of subcortical regions is also consistent with a constructive view of emotion^69^ – the perception of faces constructs a multi-modal explanation of the sensory stimuli and context, triggering an emotion reflected in the instance of emotion depicted in the face stimuli.

Finally, we sought to understand the underlying biology of alprazolam action during facial affect processing by evaluating correlations between drug-induced differences in control input and neurotransmitter receptor maps obtained through PET/SPECT imaging. Due to alprazolam’s known mechanism of action as a positive allosteric modulator of GABA_A_ receptors^27^, we expected drug-induced differences in control input to align spatially with GABA_A_ receptors. We found that these correlations, although trending positive, were not statistically significant. However, drug difference maps were positively correlated with serotonergic receptors and negatively correlated with dopamine receptors. These results indicate that the effect of alprazolam may manifest primarily through driving complementary serotonergic and dopaminergic neuromodulatory systems^70–72^, perhaps shedding light on a possible mechanistic basis of its well-documented sedative and anxiolytic effects. Our results align with previous studies which have shown that benzodiazepines like most drugs do not act in isolation, and their clinical effects likely result from affecting multiple interacting neurotransmitter systems^72^. Overall, our findings and approach highlight the utility of network control theory in understanding the neurobiological basis of drug action in the brain.

## Limitations

This study has a number of limitations. The first relates to the sample under study. As discussed previously^34^, the sample studied did not include patients with schizophrenia. The use of control energy measures as a schizophrenia endophenotype remains to be tested in larger samples, and control energy abnormalities found here in family members will need to be tested in patients with frank illness. Further, this particular sample of relatives did not exhibit marked emotion processing abnormalities assessed by behavioral performance (Supplementary Data Files 6), unlike previous results with larger cohorts^8^. It is possible that the lack of more pronounced differences in control energy parameters between relatives and controls reflects the relative normality of this sample. Further, we did not find significant associations between schizotypy scores and control energy measures, perhaps reflecting the lack of significant variation in clinical risk for psychosis in this sample. Second, while GLM parameter estimates provide a reliable indicator of brain activation patterns at the group level in response to task stimuli, this approach fails to account for dynamic variations in activation, including latencies in interactions among different brain regions. Recently developed network approaches could prove useful in studying the effect of drug and schizophrenia status on these dynamics^73,74^. Lastly, we were able to analyze neuroimaging data only from a limited slab that was chosen for high-resolution coverage of a specific set of emotion processing areas including fusiform and orbitofrontal cortex in addition to subcortical and limbic regions. The power of the network control approach in uncovering whole-brain network dynamics was thus limited to regions covered within the slab. Future studies could extend the network control approach to whole-brain imaging data obtained during facial affect processing.

### Conclusion

In summary, the network control approach described here is a powerful mechanistic framework to uncover endophenotypes of psychiatric disease and to investigate the effect of pharmacologic manipulation on the brain. Brain regions identified by the network control approach can be used to inform more targeted drug development for neuropsychiatric disorders, in addition to informing novel regions for stimulation through paradigms such as rTMS^75^. Further, control energy measures represent a readout of brain function and can be used to investigate abnormalities in various cognitive domains such as working memory^43^ and sensorimotor function^76^.

## Code availability

All analysis code is available at https://github.com/arunsm/alpraz-project.git

## Acknowledgements

This original study in which this data was collected was funded by AstraZeneca Pharmaceuticals LP. EJC acknowledges support from the National Institute of Mental Health (F30 MH118871). LP acknowledges support from NIH R01MH113550. BL acknowledges support from T32MH014654. The authors wish to acknowledge Jeff Valdez and Eve Overton for data acquisition and Mark A. Elliott for MR methods development.

## Citation Diversity Statement

Recent work in several fields of science has identified a bias in citation practices such that papers from women and other minority scholars are under-cited relative to the number of such papers in the field^77–81^. Here we sought to proactively consider choosing references that reflect the diversity of the field in thought, form of contribution, gender, race, ethnicity, and other factors. First, we obtained the predicted gender of the first and last author of each reference by using databases that store the probability of a first name being carried by a woman^81,82^. By this measure (and excluding self-citations to the first and last authors of our current paper), our references contain 7.23% woman(first)/woman(last), 12.66% man/woman, 22.09% woman/man, and 58.01% man/man. This method is limited in that a) names, pronouns, and social media profiles used to construct the databases may not, in every case, be indicative of gender identity and b) it cannot account for intersex, non-binary, or transgender people. Second, we obtained predicted racial/ethnic category of the first and last author of each reference by databases that store the probability of a first and last name being carried by an author of color^83,84^. By this measure (and excluding self-citations), our references contain 12.63% author of color (first)/author of color(last), 13.13% white author/author of color, 19.37% author of color/white author, and 54.87% white author/white author. This method is limited in that a) names and Florida Voter Data to make the predictions may not be indicative of racial/ethnic identity, and b) it cannot account for Indigenous and mixed-race authors, or those who may face differential biases due to the ambiguous racialization or ethnicization of their names. We look forward to future work that could help us to better understand how to support equitable practices in science.

## References

1. Heimberg, C., Gur, R. E., Erwin, R. J., Shtasel, D. L. & Gur, R. C. Facial emotion discrimination: III. Behavioral findings in schizophrenia. Psychiatry Res. 42, 253–265 (1992).

2. Kohler, C. G., Walker, J. B., Martin, E. A., Healey, K. M. & Moberg, P. J. Facial emotion perception in schizophrenia: A meta-analytic review. Schizophr. Bull. 36, 1009–1019 (2010).

3. Penn, D. L., Spaulding, W., Reed, D. & Sullivan, M. The relationship of social cognition to ward behavior in chronic schizophrenia. Schizophr. Res. 20, 327–335 (1996).

4. Couture, S. M., Penn, D. L. & Roberts, D. L. The functional significance of social cognition in schizophrenia: A review. Schizophr. Bull. 32, (2006).

5. Phillips, L. K. & Seidman, L. J. Emotion processing in persons at risk for schizophrenia. Schizophr. Bull. 34, 888–903 (2008).

6. Gur, R. E. et al. Neurocognitive endophenotypes in a multiplex multigenerational family study of schizophrenia. Am. J. Psychiatry 164, 813–819 (2007).

7. Allott, K. A. et al. Emotion recognition in unaffected first-degree relatives of individuals with first-episode schizophrenia. Schizophr. Res. 161, 322–328 (2015).

8. Martin, D. et al. Systematic review and meta-analysis of the relationship between genetic risk for schizophrenia and facial emotion recognition. Schizophr. Res. 218, 7–13 (2020).

9. Rasetti, R. et al. Evidence that altered amygdala activity in schizophrenia is related to clinical state and not genetic risk. Am. J. Psychiatry 166, 216–225 (2009).

10. Gur, R. E. et al. Limbic activation associated with misidentification of fearful faces and flat affect in schizophrenia. Arch. Gen. Psychiatry 64, 1356–1366 (2007).

11. Satterthwaite, T. D. et al. Association of enhanced limbic response to threat with decreased cortical facial recognition memory response in schizophrenia. Am. J. Psychiatry 167, 418– 426 (2010).

12. Anticevic, A. et al. Amygdala recruitment in schizophrenia in response to aversive emotional material: A meta-analysis of neuroimaging studies. Schizophr. Bull. 38, 608–621 (2012).

13. Butler, P. D. et al. Sensory contributions to impaired emotion processing in schizophrenia. Schizophr. Bull. 35, 1095–1107 (2009).

14. Vuilleumier, P. & Pourtois, G. Distributed and interactive brain mechanisms during emotion face perception: Evidence from functional neuroimaging. Neuropsychologia 45, 174–194 (2007).

15. Satterthwaite, T. D. et al. Opposing amygdala and ventral striatum connectivity during emotion identification. Brain Cogn. 76, 353–363 (2011).

16. Dolcos, F. et al. Emerging directions in emotional episodic memory. Front. Psychol. 8, 1– 25 (2017).

17. Li, H., Chan, R. C. K., McAlonan, G. M. & Gong, Q. Y. Facial emotion processing in schizophrenia: A meta-analysis of functional neuroimaging data. Schizophr. Bull. 36, 1029– 1039 (2010).

18. Park, H. Y. et al. Decreased neural response for facial emotion processing in subjects with high genetic load for schizophrenia. Prog. Neuro-Psychopharmacology Biol. Psychiatry 71, 90–96 (2016).

19. Taylor, S. F. & Tso, I. F. GABA abnormalities in schizophrenia: a methodological review of in vivo studies. Schizophr. Res. 167, 84–90 (2015).

20. Hashimoto, T. et al. Alterations in GABA-related transcriptome in the dorsolateral prefrontal cortex of subjects with schizophrenia. Mol. Psychiatry 13, 147–161 (2008).

21. Hashimoto, T. et al. Conserved Regional Patterns of GABA-Related Transcript Expression in the Neocortex of Subjects With Schizophrenia. Am. J. Psychiatry 165, 479–489 (2008).

22. Yoon, J. H. et al. GABA concentration is reduced in visual cortex in schizophrenia and correlates with orientation-specific surround suppression. J. Neurosci. 30, 3777–3781 (2010).

23. Spokes, E. G. S., Garrett, N. J., Rossor, M. N. & Iversen, L. L. Distribution of GABA in post-mortem brain tissue from control, psychotic and Huntington’s chorea subjects. J. Neurol. Sci. 48, 303–313 (1980).

24. Simpson, M. D. C., Slater, P., Deakin, J. F. W., Royston, M. C. & Skan, W. J. Reduced GABA uptake sites in the temporal lobe in schizophrenia. Neurosci. Lett. 107, 211–215 (1989).

25. Wise, R. G. & Tracey, I. The role of fMRI in drug discovery. J. Magn. Reson. Imaging 23, 862–876 (2006).

26. Davis, M. The Role Of The Amygdala In Fear And Anxiety. Annu. Rev. Neurosci. 15, 353– 375 (1992).

27. Verster, J. C. & Volkerts, E. R. Clinical Pharmacology, Clinical Efficacy, and Behavioral Toxicity of Alprazolam: A Review of the Literature. CNS Drug Rev. 10, 45–76 (2004).

28. Ashton, H. The diagnosis and management of benzodiazepine dependence. Curr. Opin. Psychiatry 18, 249–255 (2005).

29. Buchanan, T. W., Karafin, M. S. & Adolphs, R. Selective effects of Triazolam on memory for emotional, relative to neutral, stimuli: Differential effects on gist versus detail. Behav. Neurosci. 117, 517–525 (2003).

30. Coupland, N. J., Singh, A. J., Sustrik, R. A., Ting, P. & Blair, R. J. Effects of diazepam on facial emotion recognition. J. Psychiatry Neurosci. 28, 452–463 (2003).

31. Garcez, H. et al. Effects of benzodiazepines administration on identification of facial expressions of emotion: a meta-analysis. Psychopharmacology (Berl). 237, (2020).

32. Paulus, M. P., Feinstein, J. S., Castillo, G., Simmons, A. N. & Stein, M. B. Dose-dependent decrease of activation in bilateral amygdala and insula by lorazepam during emotion processing. Arch. Gen. Psychiatry 62, 282–288 (2005).

33. Del-Ben, C. M. et al. Effects of diazepam on BOLD activation during the processing of aversive faces. J. Psychopharmacol. 26, 443–451 (2012).

34. Wolf, D. H. et al. Amygdala abnormalities in first-degree relatives of individuals with schizophrenia unmasked by benzodiazepine challenge. Psychopharmacology (Berl). 218, 503–512 (2011).

35. Liu, Y. Y., Slotine, J. J. & Barabási, A. L. Controllability of complex networks. Nature 473, 167–173 (2011).

36. Pasqualetti, F., Zampieri, S. & Bullo, F. Controllability metrics, limitations and algorithms for complex networks. IEEE Trans. Control Netw. Syst. 1, 40–52 (2014).

37. Bassett, D. S. & Sporns, O. Network neuroscience. Nat. Neurosci. 20, 353–364 (2017).

38. Yan, G. et al. Network control principles predict neuron function in the Caenorhabditis elegans connectome. Nature 550, 519–523 (2017).

39. Kim, J. Z. et al. Role of graph architecture in controlling dynamical networks with applications to neural systems. Nat. Phys. 14, 91–98 (2018).

40. Brynildsen, J. K. et al. Gene coexpression patterns predict opiate-induced brain-state transitions. Proc. Natl. Acad. Sci. U. S. A. 117, 19556–19565 (2020).

41. Szymula, K. P., Pasqualetti, F., Graybiel, A. M., Desrochers, T. M. & Bassett, D. S. Habit learning supported by efficiently controlled network dynamics in naive macaque monkeys. arXiv XXX, (2020).

42. Gu, S. et al. Controllability of structural brain networks. Nat. Commun. 6, 1–10 (2015).

43. Braun, U. et al. Brain state stability during working memory is explained by network control theory, modulated by dopamine D1/D2 receptor function, and diminished in schizophrenia. bioRxiv 679670 (2019) doi:10.1101/679670.

44. Lee, W. H., Rodrigue, A., Glahn, D. C., Bassett, D. S. & Frangou, S. Heritability and Cognitive Relevance of Structural Brain Controllability. Cereb. Cortex 30, 3044–3054 (2020).

45. Bernhardt, B. C. et al. Temporal lobe epilepsy: Hippocampal pathology modulates connectome topology and controllability. Neurology 92, E2209–E2220 (2019).

46. Jeganathan, J. et al. Fronto-limbic dysconnectivity leads to impaired brain network controllability in young people with bipolar disorder and those at high genetic risk. NeuroImage Clin. 19, 71–81 (2018).

47. Karrer, T. M. et al. A practical guide to methodological considerations in the controllability of structural brain networks. J. Neural Eng. 17, 26031 (2020).

48. Betzel, R. F., Gu, S., Medaglia, J. D., Pasqualetti, F. & Bassett, D. S. Optimally controlling the human connectome: The role of network topology. Sci. Rep. 6, 1–14 (2016).

49. Gu, S. et al. Optimal trajectories of brain state transitions. Neuroimage 148, 305–317 (2017).

50. Stiso, J. et al. White Matter Network Architecture Guides Direct Electrical Stimulation through Optimal State Transitions. Cell Rep. 28, 2554-2566.e7 (2019).

51. Cornblath, E. J. et al. Temporal sequences of brain activity at rest are constrained by white matter structure and modulated by cognitive demands. Commun. Biol. 3, 1–12 (2020).

52. Gur, R. C. et al. A method for obtaining 3-dimensional facial expressions and its standardization for use in neurocognitive studies. J. Neurosci. Methods 115, 137–143 (2002).

53. Esteban, O. et al. fMRIPrep: a robust preprocessing pipeline for functional MRI. Nat. Methods 16, 111–116 (2019).

54. Woolrich, M. W., Ripley, B. D., Brady, M. & Smith, S. M. Temporal autocorrelation in univariate linear modeling of FMRI data. Neuroimage 14, 1370–1386 (2001).

55. Ciric, R. et al. Mitigating head motion artifact in functional connectivity MRI. Nat. Protoc. 13, 2801–2826 (2018).

56. Loughead, J., Gur, R. C., Elliott, M. & Gur, R. E. Neural circuitry for accurate identification of facial emotions. Brain Res. 1194, 37–44 (2008).

57. Cammoun, L. et al. Mapping the human connectome at multiple scales with diffusion spectrum MRI. J. Neurosci. Methods 203, 386–397 (2012).

58. Dukart, J. et al. JuSpace: A tool for spatial correlation analyses of magnetic resonance imaging data with nuclear imaging derived neurotransmitter maps. Hum. Brain Mapp. 1– 12 (2020) doi:10.1002/hbm.25244.

59. Dukart, J. et al. Cerebral blood flow predicts differential neurotransmitter activity. Sci. Rep. 8, 1–11 (2018).

60. Alakurtti, K. et al. Long-term test-retest reliability of striatal and extrastriatal dopamine D2/3 receptor binding: Study with [11C]raclopride and high-resolution PET. J. Cereb. Blood Flow Metab. 35, 1199–1205 (2015).

61. Hesse, S. et al. Central noradrenaline transporter availability in highly obese, non-depressed individuals. Eur. J. Nucl. Med. Mol. Imaging 44, 1056–1064 (2017).

62. Kaller, S. et al. Test–retest measurements of dopamine D1-type receptors using simultaneous PET/MRI imaging. Eur. J. Nucl. Med. Mol. Imaging 44, 1025–1032 (2017).

63. Gómez, F. J. G., Huertas, I., Ram\’\irez, J. A. L. & Sol\’\is, D. G. Elaboración de una plantilla de SPM para la normalización de imágenes de PET con 18F-DOPA. Imagen Diagnóstica 9, 23–25 (2018).

64. Savli, M. et al. Normative database of the serotonergic system in healthy subjects using multi-tracer PET. Neuroimage 63, 447–459 (2012).

65. Benjamini, Y. & Hochberg, Y. Controlling the False Discovery Rate: A Practical and Powerful Approach to Multiple Testing. Journal of the Royal Statistical Society. Series B (Methodological) vol. 57 289–300 (1995).

66. Tsao, D. Y., Moeller, S. & Freiwald, W. A. Comparing face patch systems in macaques and humans. Proc. Natl. Acad. Sci. U. S. A. 105, 19514–19519 (2008).

67. Saygin, Z. M. et al. Anatomical connectivity patterns predict face selectivity in the fusiform gyrus. Nat. Neurosci. 15, 321–327 (2012).

68. Adolphs, R., Tranel, D., Damasio, H. & Damasio, A. R. Fear and the human amygdala. J. Neurosci. 15, 5879–5891 (1995).

69. Barrett, L. F. & Russell, J. A. The psychological construction of emotion. (Guilford Publications, 2014).

70. Giardino, L., Zanni, M., Pozza, M., Bettelli, C. & Covelli, V. Dopamine receptors in the striatum of rats exposed to repeated restraint stress and alprazolam treatment. Eur. J. Pharmacol. 344, 143–147 (1998).

71. Bentue-Ferrer, D. et al. Role of dopaminergic and serotonergic systems on behavioral stimulatory effects of low-dose alprazolam and lorazepam. Eur. Neuropsychopharmacol. 11, 41–50 (2001).

72. Avery, M. C. & Krichmar, J. L. Neuromodulatory systems and their interactions: A review of models, theories, and experiments. Front. Neural Circuits 11, 1–18 (2017).

73. Goghari, V. M., Sanford, N., Spilka, M. J. & Woodward, T. S. Task-Related Functional Connectivity Analysis of Emotion Discrimination in a Family Study of Schizophrenia. Schizophr. Bull. 43, 1348–1362 (2017).

74. Lavigne, K. M., Menon, M. & Woodward, T. S. Functional Brain Networks Underlying Evidence Integration and Delusions in Schizophrenia. Schizophr. Bull. 46, 175–183 (2019).

75. Balderston, N. L. et al. Low-frequency parietal repetitive transcranial magnetic stimulation reduces fear and anxiety. Transl. Psychiatry (2020) doi:10.1038/s41398-020-0751-8.

76. Stiso, J. et al. Learning in brain-computer interface control evidenced by joint decomposition of brain and behavior. J. Neural Eng. (2020).

77. Mitchell, S. M., Lange, S. & Brus, H. Gendered Citation Patterns in International Relations Journals. Int. Stud. Perspect. 14, 485–492 (2013).

78. Maliniak, D., Powers, R. & Walter, B. F. The gender citation gap in international relations. International Organization vol. 67 (2013).

79. Caplar, N., Tacchella, S. & Birrer, S. Quantitative evaluation of gender bias in astronomical publications from citation counts. Nat. Astron. 1, (2017).

80. Dion, M. L., Sumner, J. L. & Mitchell, S. M. L. Gendered Citation Patterns across Political Science and Social Science Methodology Fields. Polit. Anal. 26, 312–327 (2018).

81. Dworkin, J. D. et al. The extent and drivers of gender imbalance in neuroscience reference lists. Nat. Neurosci. 23, 918–926 (2020).

82. Zhou, D. et al. Gender Diversity Statement and Code Notebook v1.0. (2020) doi:10.5281/zenodo.3672110.

83. Ambekar, A., Ward, C., Mohammed, J., Male, S. & Skiena, S. Name-ethnicity classification from open sources. Proc. ACM SIGKDD Int. Conf. Knowl. Discov. Data Min. 49–57 (2009) doi:10.1145/1557019.1557032.

84. Sood, G. & Laohaprapanon, S. Predicting race and ethnicity from the sequence of characters in a name. arXiv Prepr. arXiv1805.02109 (2018).

