## Supplementary Information for "Alprazolam modulates persistence energy during emotion processing in first-degree relatives of individuals with schizophrenia: a network control study"

### Supplementary Methods

#### Subject assessment

The clinical evaluation included the Diagnostic Interview for Genetic Studies (DIGS)[1] and Family Interview for Genetics Studies (FIGS)[2]. Schizotypy was assessed with the Structured Interview for Schizotypy (SIS)[3] included in the DIGS; total SIS scores were calculated as the sum of global items (SIS scores were unavailable for 1 family member). Anxiety was assessed using the State Trait Anxiety Inventory (STAI)[4]; state anxiety was measured three times at each study session: prior to drug administration (baseline), just before scanning, and immediately after the scanning session. Neurocognitive function was assessed with the Penn Computerized Neurocognitive Battery[5–7]. Interviews were administered by trained assessors with demonstrated reliability on symptom scales (reliability criterion: intraclass correlation coefficient >0.90).

Participants had no current Axis I psychiatric disorders, or any history of a psychotic disorder. Axis II disorders were also exclusionary, with the exception of Cluster A personality disorders for family member participants, as Cluster A personality traits might reflect genetic risk for schizophrenia. Family members had a first-degree relative with DSM-IV schizophrenia as determined through the FIGS interview, supporting medical records, and when possible a DIGS assessment of the ill family member. Control participants had no family history of a psychotic disorder or bipolar disorder in first- or second-degree relatives. Participants had no history of alcohol or other substance use disorders within the past 6 months, no history of benzodiazepine abuse or dependence at any time, and no recent substance use by history and confirmed by urine drug screen on each study day. Use of alcohol in the 24 hours preceding the study was exclusionary, as was use of psychoactive substances or medications within the prior 2 weeks, except for nicotine and caffeine where subjects maintained regular use to avoid withdrawal effects. Participants had no history of neurological or medical disorders or injuries affecting brain function, or which would contraindicate MRI scanning or benzodiazepine administration.

#### BOLD and structural image acquisition procedures and parameters

Subjects were placed in the scanner supine. Earplugs were used to muffle scanner noise. Head fixation was ensured by a foam-rubber device mounted on the head coil. Pulse and respiration monitors were attached. Stimuli were rear-projected to the center of the visual field using a PowerLite 7300 video projector (Epson America, Inc.; Long Beach, CA) and viewed through a head coil mounted mirror. Stimulus presentation was synchronized with image acquisition using

Presentation software (Neurobehavioral Systems, Inc., Albany, CA). Subjects provided responses with a non-ferromagnetic response device (fORP, Current Designs, Inc., Philadelphia, PA) using their dominant hand. Button press responses and response latencies were recorded.

Whole-brain structural data were obtained with a 5-minute magnetization-prepared, rapid acquisition gradient-echo T1-weighted image (MPRAGE, TR 1620 ms, TE 3.87 ms, FOV 180×240 mm, matrix 192×256, effective voxel resolution of 1×1×1 mm). BOLD fMRI data was obtained as a slab single-shot gradient-echo (GE) echoplanar sequence using the following parameters: TR/TE=3000/32 ms, FOV=240 mm, matrix= 128 X 128, slice thickness/gap=2/0 mm, 30 slices, effective voxel resolution of 1.875x1.875x2 mm. Slices were obtained obliquely (axial/coronal) in order to reduce signal distortion in ventral brain regions.

**Diffusion imaging data acquisition**

High resolution anatomical brain images were collected from 10 healthy young adults (23.9 ± 3.6 years; 20-31 years; 70% female). The participants underwent a 53:24 minute diffusion spectrum imaging (DSI) scan with 730 diffusion directions (maximum $b$-value = 5010 s/mm$^2$, 21 $b$ = 0 images, TR = 4300 ms, TE = 102 ms, matrix size = 144×144, field of view = 260×260 mm$^2$, slice number = 87, resolution =1.8×1.8×1.8 mm$^3$, multi-band acceleration factor = 3). Additionally, T1-weighted images were obtained using an MPRAGE sequence (TR = 2500 ms, TE = 2.18 ms, flip angle = 7 degrees, slice number = 208, slice thickness = 0.9 mm). Both scans were acquired on a Siemens Magnetom Prisma 3 Tesla scanner with a 64-channel head coil. The DSI study was conducted separately from the BOLD study and was approved by the Institutional Review Board of the University of Pennsylvania. All participants provided informed consent in writing.

**Image processing**

Anatomical and functional BOLD fMRI images were processed using *fMRIPrep* 1.2.6[8], which is based on *Nipype* 1.1.7[9]. Image processing pipeline details are described below.

**Anatomical data preprocessing**

T1-weighted (T1w) images were corrected for intensity non-uniformity (INU) using N4BiasFieldCorrection in ANTS 2.2.0[10]. A T1w-reference map was computed after registration of T1w images (after INU-correction) using mri_robust_template[11] (FreeSurfer 6.0.1). The T1w-reference was then skull-stripped using antsBrainExtraction.sh (ANTs 2.2.0) with OASIS as the target template. Brain surfaces were reconstructed using recon-all[12] (FreeSurfer 6.0.1), and the brain mask estimated previously was refined with a custom variation of the method to reconcile ANTs-derived and FreeSurfer-derived segmentations of the cortical gray matter of Mindboggle[13]. Spatial normalization to the ICBM 152 Nonlinear Asymmetrical template version

2009c[14] was performed through nonlinear registration with antsRegistration[15] (ANTs 2.2.0), using brain-extracted versions of both the T1w volume and template. Brain tissue segmentation of cerebrospinal fluid (CSF), white matter (WM), and gray matter (GM) was performed on the brain-extracted T1w using fast[16] (FSL 5.0.9).

**Functional data preprocessing**

For each of the 4 BOLD runs per subject (across all tasks and sessions), the following preprocessing was performed. First, a reference volume and its skull-stripped version were generated using a custom methodology of *fMRIPrep*. A deformation field to correct for susceptibility distortions was estimated based on *fMRIPrep*'s *fieldmap-less* approach. The deformation field is that resulting from co-registering the BOLD reference to the same-subject T1w-reference with its intensity inverted[17,18]. Registration was performed with antsRegistration (ANTs 2.2.0), and the process regularized by constraining deformation to be nonzero only along the phase-encoding direction, and modulated with an average fieldmap template[19]. Based on the estimated susceptibility distortion, an unwarped BOLD reference was calculated for a more accurate co-registration with the anatomical reference. The BOLD reference was then co-registered to the T1w reference using bbregister (FreeSurfer) which implements boundary-based registration[20]. Co-registration was configured with nine degrees of freedom to account for distortions remaining in the BOLD reference. Head-motion parameters with respect to the BOLD reference (transformation matrices, and six corresponding rotation and translation parameters) were estimated before any spatiotemporal filtering using mcflirt[21] (FSL 5.0.9). The BOLD time-series were resampled to surfaces on the following spaces: *fsaverage5*. The BOLD time-series were resampled onto their original, native space by applying a single, composite transform to correct for head-motion and susceptibility distortions. These resampled BOLD time-series will be referred to as *preprocessed BOLD in original space*, or just *preprocessed BOLD*. The BOLD time-series were resampled to MNI152NLin2009cAsym standard space, generating a *preprocessed BOLD run in MNI152NLin2009cAsym space*. First, a reference volume and its skull-stripped version were generated using a custom methodology of *fMRIPrep*. Several confounding timeseries were calculated based on the *preprocessed BOLD*: framewise displacement (FD), DVARS, and three region-wise global signals. FD and DVARS were calculated for each functional run, both using their implementations in *Nipype* (following the definitions in Ref. [22]). The three global signals were extracted within the CSF, the WM, and the whole-brain masks. Additionally, a set of physiological regressors were extracted to allow for component-based noise correction (*CompCor*)[23]. Principal components were estimated after high-pass filtering the *preprocessed BOLD* time-series (using a discrete cosine filter with 128s cut-off) for the two *CompCor* variants: temporal (tCompCor) and anatomical (aCompCor). Six tCompCor components were then calculated from the top 5% variable voxels within a mask covering the subcortical regions. This subcortical mask was obtained by heavily eroding the brain mask, which ensures it does not include cortical grey matter regions. For aCompCor, six components were calculated within the intersection of the aforementioned mask and the union of CSF and WM masks calculated in T1w space, after their projection to the native space of each functional run (using the inverse BOLD-to-T1w transformation). The head-motion estimates calculated in the correction step were also placed within the corresponding confounds file. All resamplings were performed with *a single interpolation step* by composing all the pertinent transformations (i.e. head-motion transform matrices, susceptibility distortion correction when available, and co-registrations to anatomical and template spaces). Gridded (volumetric) resamplings were performed using antsApplyTransforms (ANTs), configured with Lanczos interpolation to minimize the smoothing effects of other kernels[24]. Non-gridded (surface) resamplings were performed using mri_vol2surf (FreeSurfer). Many internal operations of *fMRIPrep* use *Nilearn* 0.5.0[25], mostly within the functional processing workflow. For more details of the pipeline, see the section corresponding to workflows in *fMRIPrep*'s documentation.

### Diffusion imaging data preprocessing

As described in Ref. [26], individual DSI scans were skull-stripped, realigned, and motion-corrected using an improved average $b$=0 reference image. The preprocessing was implemented in Nipype[9] using the Advanced Normalization Tools (ANTs)[15] for image registration. We quantified the diffusion at different orientations in each voxel using the generalized q-sampling reconstruction method[27] in DSI Studio (dsi-studio.labsolver.org). Based on the derived quantitative anisotropy values, we performed deterministic tractography across the whole-brain[28]. For each participant, we generated 1,000,000 streamlines with a maximum length of 500 mm[29] and a maximum turning angle of 35 degrees[30].

### Construction of structural brain networks

From the diffusion imaging data, we constructed a structural brain network for each participant. Consistent with previous work[26,31,32], we defined nodes of the network as brain regions according to the 234-node Lausanne atlas[33]. For this purpose, the Lausanne parcels were dilated by 4 mm so that the parcels reached down into the white matter enough to ensure accurate sampling of underlying fibers. In the process of dilation, some voxels were assigned to two or more regions of interest; to eradicate this redundancy, we assigned each voxel to the mode of its neighbors[34]. After warping the parcellation into the subject's diffusion space, we weighted each edge in the network by the quantitative anisotropy (QA) for each pair of brain regions, corrected for their volume. The brain stem parcel was excluded due to its hyperconnectivity with other nodes in the network. Overall, we constructed a 233×233 sparse, weighted, and undirected adjacency matrix for each participant.

### Alternate analyses

Alternate analyses were conducted to supplement the results presented in Figure 2 in the main text, by replacing categorical group and drug indicators with continuous variables. When alprazolam blood levels were used in place of categorical drug indicator in the mixed model, we saw significant main effects of group ($\gamma_{01}$=-0.045, p=0.014, df=74) and drug ($\beta_{1i}$=-0.004, p=0.048, df=74), with a sub-threshold trend toward a group×drug interaction ($\gamma_{11}$=0.005, p=0.064, df=74) during threat emotion identification. No effects were found during identification of non-threat and neutral categories. In the emotion memory task, results remained similar to using categorical variables (Supplementary Data Files 2).

Next, we used linear mixed models to assess relations between persistence energy and psychopathology by replacing the categorical group indicator with the total score on the structured interview for schizotypy (SIS). Here, we did not observe any significant effects of drug or SIS except for a drug main effect ($\beta_{1i}$=0.055, p=0.031, df=79) and a trend towards a drug×SIS interaction ($\gamma_{11}$=-0.003, p=0.071, df=79) during neutral emotion identification (Supplementary Data Files 3).

### Null model analysis

To elucidate the differential contribution of structural brain networks and brain activation maps on control energy measures, we performed analyses using a series of null models. Results from these analyses are described below.

### Null models of structural brain networks

To study the impact of structural brain networks on persistence energy, we re-computed all control energies using 500 randomized null models of structural brain networks that preserved the degree distribution. Random brain networks were generated using the randmio_und function implemented in the Brain Connectivity Toolbox (https://sites.google.com/site/bctnet/). Using the control energies obtained from the null models, we refit the linear mixed effects models used to generate results for Figure 2 in the main text and evaluated the significance of drug, group and group×drug interactions. Results from this analysis are shown in Figure S4 (see also Supplementary Data Files 8). Coefficients for group×drug interactions during threat emotion memory reached significance ($p < 0.05$) in 1.8% (9/500) of structural null model iterations. None of the other coefficients reached a level of significance ($p < 0.05$) in any of the null model iterations. As predicted, these results show that the differential effect of alprazolam on persistence energy in relatives and controls during recall of threatening faces is partially driven by the precise topology of the underlying structural brain networks.

**Null models of brain activation patterns**

To study the impact of spatial distribution of activity patterns on control properties, we re-computed persistence energy after spatially randomizing beta maps from GLMs[35]. Using the persistence energy obtained from 500 iterations of spatially randomized maps, we re-fit the linear mixed effects models and evaluated the significance of coefficients for drug, group and group×drug interaction as above. Results from this analysis are shown in Figure S5 (see also Supplementary Data Files 9). Main effect of group reached a level of significance ($p < 0.05$) in 0.02% (1/500) of iterations for threat emotion identification and 0.04% (2/500) of iterations for neutral emotion identification. Mixed model coefficients for group, drug, or group×drug interaction did not reach a level of significance in any iterations for non-threat emotion identification and all the emotion memory tasks. These results show that the differential effect of alprazolam on persistence energy in relatives and controls during recall of threatening faces is strongly driven by the specific task-driven brain activation pattern.

### Supplementary Figures

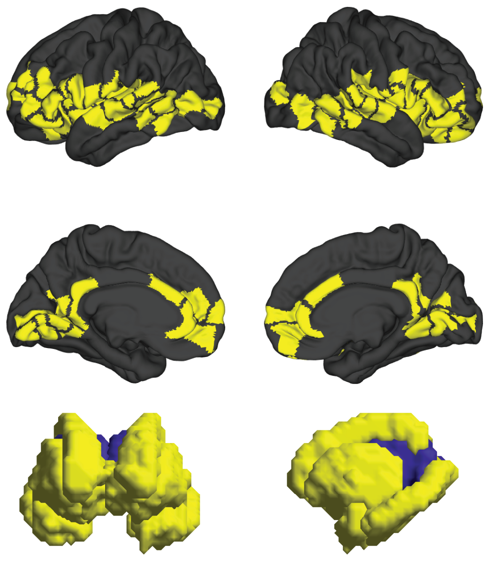

**Figure S1. Imaging slab.** BOLD images were acquired in a slab covering ventral cortical regions involved in affective processing along with subcortical areas. Lausanne parcels which have at least 50% coverage in the imaging slab are highlighted in yellow. An average mask comprising all subject masks generated during the emotion identification task was used to generate this visualization.

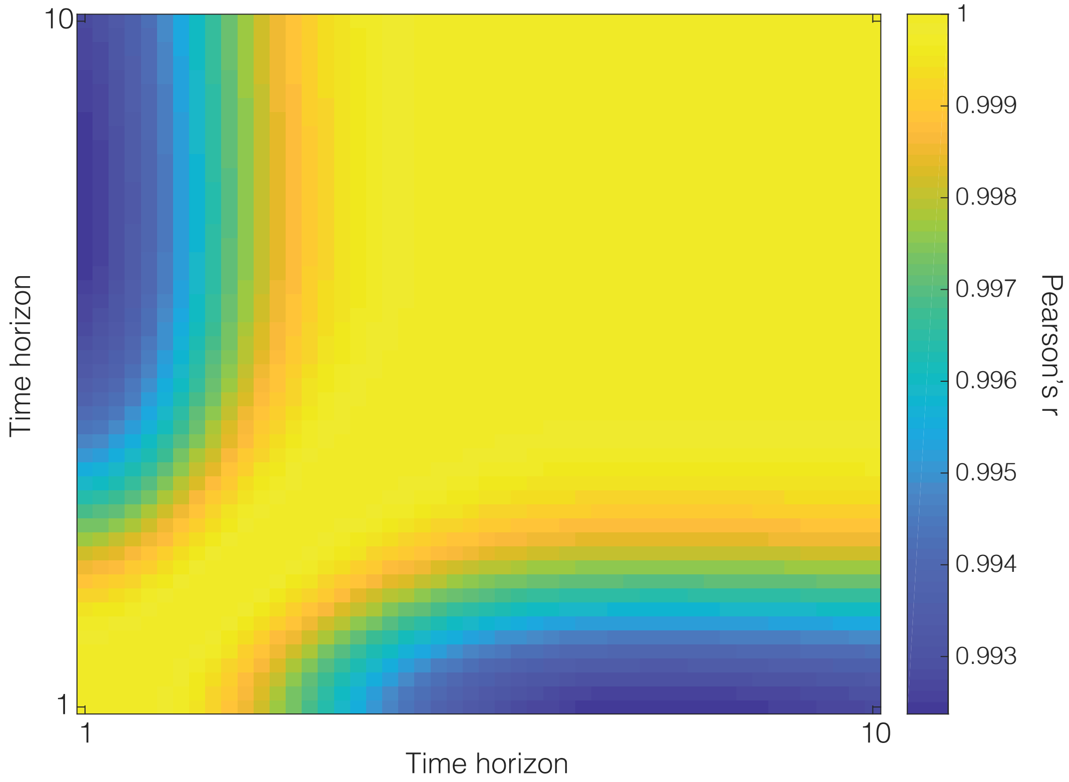


**Figure S2. Consistency of persistence energy during threat emotion identification for different choices of time horizon.** The heat map shows the correlation matrix of different time horizons, with each entry corresponding to the Pearson correlation coefficient between persistence energy for all subjects based on two different time horizon choices. Note that the Pearson correlation coefficients are quite high, indicating that the choice of time horizon did not significantly affect the relative estimated values of minimal control energy.

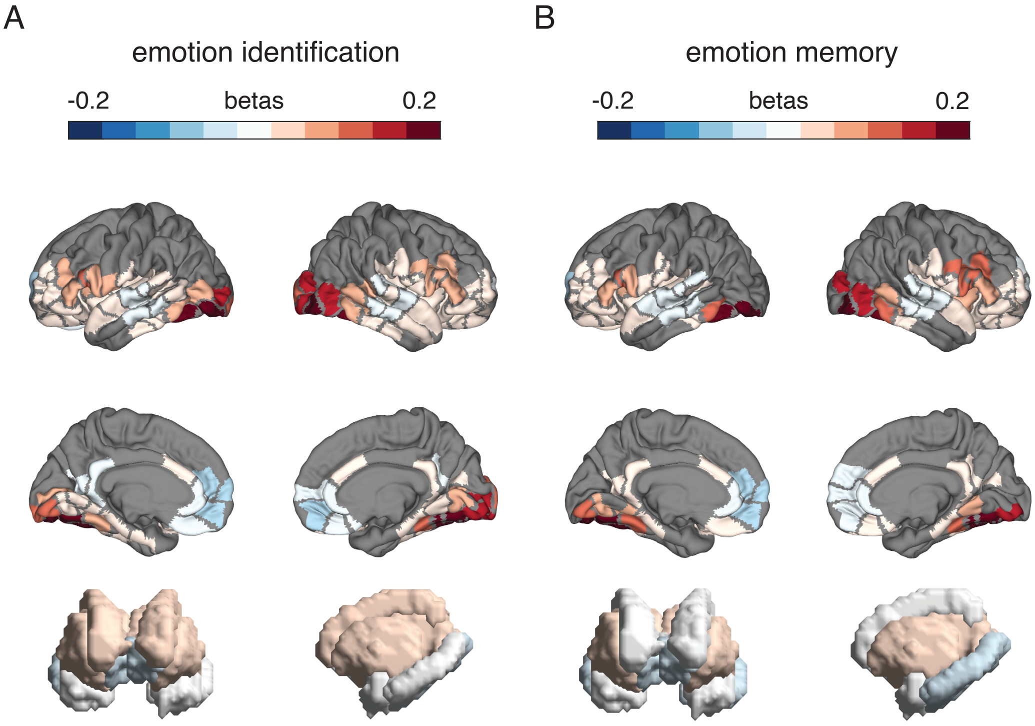


**Figure S3. Average spatial maps of normalized GLM betas.** (A) Average spatial maps of GLM betas (normalized by Euclidean norm) for threat emotion identification, shown on surface renderings of cortical and sub-cortical areas. (B) Average spatial maps of GLM betas for emotion memory, shown on surface renderings of cortical and sub-cortical areas. Areas outside the imaging slab are colored grey.

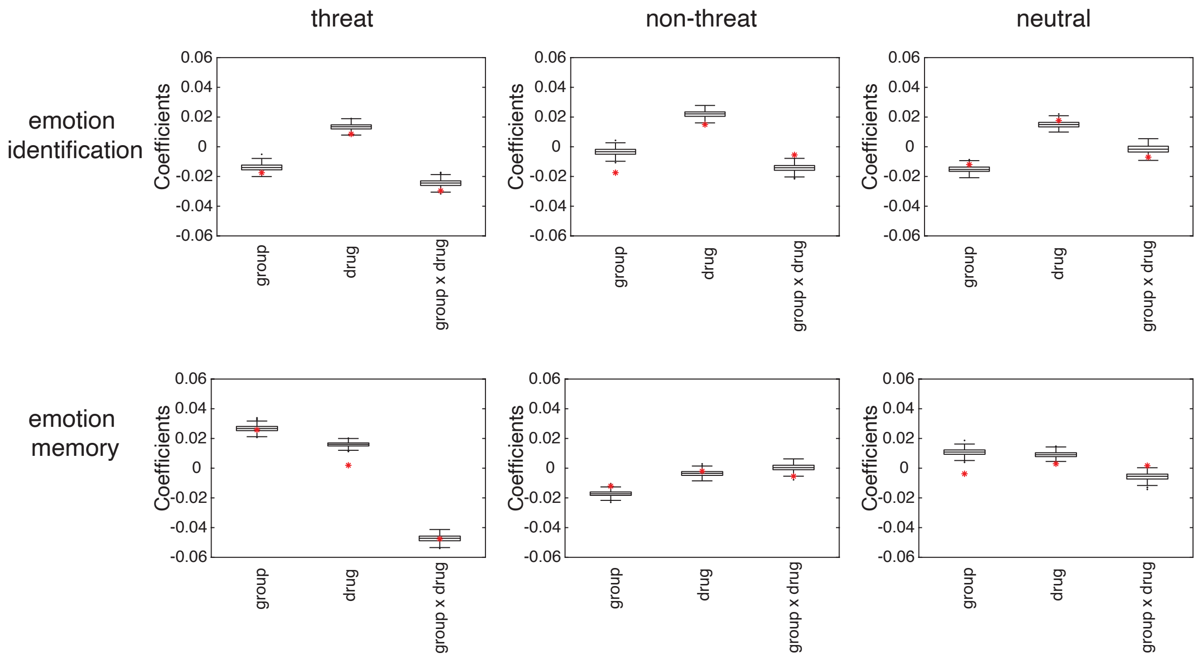


**Figure S4. Results from null models of structural brain networks.** Boxplots of group, drug, and group×drug interaction mixed model coefficients from 500 iterations of structural null models. Coefficients with original model computed with subject average structural matrix shown as red asterisks.

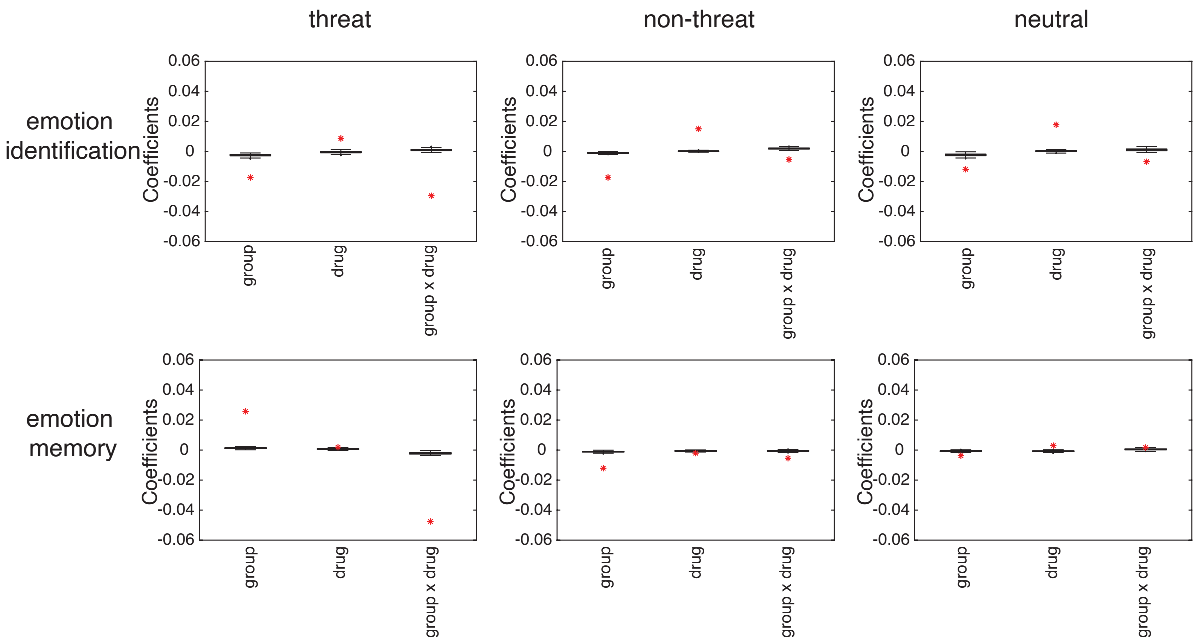


**Figure S5. Results from null models of brain activation patterns.** Boxplots of group, drug, and group×drug interaction mixed model coefficients from 500 iterations of spatial null models. Coefficients with original model computed with subject average structural matrix shown as red asterisks.

### Supplementary Data Files

#### Supplementary Data Files 1

Linear mixed model coefficients and statistics for drug and group effects on persistence energy during emotion identification and memory (corresponding to Eq. 1 in the main text), with drug and group indicators treated as categorical variables.

#### Supplementary Data Files 2

Linear mixed model coefficients and statistics for drug and group effects on persistence energy during emotion identification and memory (corresponding to Eq. 1 in the main text), using alprazolam blood levels as the drug indicator and treating group as a categorical factor.

#### Supplementary Data Files 3

Linear mixed model coefficients and statistics for drug and group effects on persistence energy during emotion identification and memory (corresponding to Eq. 1 in the main text), with schizotypy score (measured as the total score on the structured interview for schizotypy, SISTOTAL) used in place of the categorical group indicator.

#### Supplementary Data Files 4

Sorted control impact values for 233 Lausanne parcels during emotion identification and memory. Values for parcels outside of the imaging slab are marked as NaN.

#### Supplementary Data Files 5

Sorted beta weights from general linear models (normalized by Euclidean norm) for 233 Lausanne parcels during emotion identification and memory. Values for parcels outside of the imaging slab are marked as NaN.

#### Supplementary Data Files 6

Linear mixed model coefficients and statistics for effects of drug, group, and persistence energy on task efficiency during emotion identification and memory (corresponding to Eq. 2 in the main text).

#### Supplementary Data Files 7

Sorted regional differences in control input between alprazolam and placebo conditions for 233 Lausanne parcels during emotion identification and memory. Values for parcels outside of the imaging slab are marked as NaN.

#### Supplementary Data Files 8

Linear mixed model coefficients (corresponding to Equations 1-3 in the main text) fit against persistence energy computed using 500 iterations of structural null models, along with fraction of iterations where observed coefficients were statistically significant ($p < 0.05$).

**Supplementary Data Files 9**

Linear mixed model coefficients (corresponding to Equations 1-3 in the main text) fit against persistence energy computed using 500 iterations of spatial null models, along with fraction of iterations where observed coefficients were statistically significant ($p < 0.05$).

**Supplementary References**


1. Nurnberger, J. I. J. *et al.* Diagnostic interview for genetic studies. Rationale, unique features, and training. NIMH Genetics Initiative. *Arch. Gen. Psychiatry* **51**, 844–849 (1994).
2. Maxwell, M. E. Family Interview for Genetic Studies (FIGS): a manual for FIGS. *Bethesda, MD Clin. Neurogenetics Branch, Intramural Res. Program, Natl. Inst. Ment. Heal.* (1992).
3. Kendler, K. S., Lieberman, J. A. & Walsh, D. The Structured Interview for Schizotypy (SIS): a preliminary report. *Schizophr. Bull.* **15**, 559–571 (1989).
4. Spielberger, C. D., Gorsuch, R. L., Lushene, R., Vagg, P. R. & Jacobs, G. A. Manual for the state-trait anxiety inventory Consulting Psychologists Press. *Palo Alto, CA* (1983).
5. Gur, R. C. *et al.* Computerized neurocognitive scanning: I. Methodology and validation in healthy people. *Neuropsychopharmacology* **25**, 766–776 (2001).
6. Gur, R. C. *et al.* Computerized neurocognitive scanning: II. The profile of schizophrenia. *Neuropsychopharmacology* **25**, 777–788 (2001).
7. Gur, R. C. *et al.* A cognitive neuroscience-based computerized battery for efficient measurement of individual differences: standardization and initial construct validation. *J. Neurosci. Methods* **187**, 254–262 (2010).
8. Esteban, O. *et al.* fMRIPrep: a robust preprocessing pipeline for functional MRI. *Nat. Methods* **16**, 111–116 (2019).
9. Gorgolewski, K. *et al.* Nipype: A flexible, lightweight and extensible neuroimaging data processing framework in Python. *Front. Neuroinform.* **5**, (2011).
10. Tustison, N. J. *et al.* N4ITK: Improved N3 bias correction. *IEEE Trans. Med. Imaging* **29**, 1310–1320 (2010).
11. Reuter, M., Rosas, H. D. & Fischl, B. Highly accurate inverse consistent registration: A robust approach. *Neuroimage* **53**, 1181–1196 (2010).
12. Dale, A. M., Fischl, B. & Sereno, M. I. Cortical Surface-Based Analysis. *Neuroimage* **9**, 179–194 (1999).
13. Klein, A. et al. Mindboggling morphometry of human brains. PLoS Computational Biology vol. 13 (2017).

14. Fonov, V., Evans, A., McKinstry, R., Almli, C. & Collins, D. Unbiased nonlinear average age-appropriate brain templates from birth to adulthood. *Neuroimage* **47**, S102 (2009).
15. Avants, B. B., Epstein, C. L., Grossman, M. & Gee, J. C. Symmetric diffeomorphic image registration with cross-correlation: Evaluating automated labeling of elderly and neurodegenerative brain. *Med. Image Anal.* **12**, 26–41 (2008).
16. Zhang, Y., Brady, M. & Smith, S. Segmentation of brain MR images through a hidden Markov random field model and the expectation-maximization algorithm. *IEEE Trans. Med. Imaging* **20**, 45–57 (2001).
17. Huntenburg, J. M., Gorgolewski, K. J., Anwander, A. & Margulies, D. S. Evaluating nonlinear coregistration of BOLD EPI and T1 images. *20th Annu. Meet. Organ. Hum. Brain Mapp.* 1 (2014).
18. Wang, S. *et al.* Evaluation of field map and nonlinear registration methods for correction of susceptibility artifacts in diffusion MRI. *Front. Neuroinform.* **11**, 1–9 (2017).
19. Treiber, J. M. *et al.* Characterization and correction of geometric distortions in 814 Diffusion Weighted Images. *PLoS One* **11**, 1–9 (2016).
20. Greve, D. N. & Fischl, B. Accurate and robust brain image alignment using boundary-based registration. *Neuroimage* **48**, 63–72 (2009).
21. Jenkinson, M., Bannister, P., Brady, M. & Smith, S. Improved Optimization for the Robust and Accurate Linear Registration and Motion Correction of Brain Images. *Neuroimage* **17**, 825–841 (2002).
22. Power, J. D. *et al.* Methods to detect, characterize, and remove motion artifact in resting state fMRI. *Neuroimage* **84**, 320–341 (2014).
23. Behzadi, Y., Restom, K., Liau, J. & Liu, T. T. A component based noise correction method (CompCor) for BOLD and perfusion based fMRI. *Neuroimage* **37**, 90–101 (2007).
24. Lanczos, C. Evaluation of Noisy Data. *J. Soc. Ind. Appl. Math. Ser. B Numer. Anal.* **1**, 76–85 (1964).
25. Abraham, A. *et al.* Machine learning for neuroimaging with scikit-learn. *Front. Neuroinform.* **8**, 1–10 (2014).
26. Karrer, T. M. *et al.* A practical guide to methodological considerations in the controllability of structural brain networks. *J. Neural Eng.* **17**, 26031 (2020).
27. Yeh, F., Wedeen, V. J. & Tseng, W. I. Generalized q -Sampling Imaging. **29**, 1626–1635 (2010).
28. Yeh, F. C., Verstynen, T. D., Wang, Y., Fernández-Miranda, J. C. & Tseng, W. Y. I. Deterministic diffusion fiber tracking improved by quantitative anisotropy. *PLoS One* **8**, 1–16 (2013).
29. Cieslak, M. & Grafton, S. T. Local termination pattern analysis: A tool for comparing white matter morphology. *Brain Imaging Behav.* **8**, 292–299 (2014).
30. Bassett, D. S., Brown, J. A., Deshpande, V., Carlson, J. M. & Grafton, S. T. Conserved and variable architecture of human white matter connectivity. *Neuroimage* **54**, 1262–1279 (2011).

31. Gu, S. *et al.* Optimal trajectories of brain state transitions. *Neuroimage* **148**, 305–317 (2017).
32. Betzel, R. F., Gu, S., Medaglia, J. D., Pasqualetti, F. & Bassett, D. S. Optimally controlling the human connectome: The role of network topology. *Sci. Rep.* **6**, 1–14 (2016).
33. Cammoun, L. *et al.* Mapping the human connectome at multiple scales with diffusion spectrum MRI. *J. Neurosci. Methods* **203**, 386–397 (2012).
34. Daducci, A. *et al.* The Connectome Mapper: An Open-Source Processing Pipeline to Map Connectomes with MRI. *PLoS One* **7**, (2012).
35. Alexander-Bloch, A. F. *et al.* On testing for spatial correspondence between maps of human brain structure and function. *Neuroimage* **178**, 540–551 (2018).
